# DCAF: Unifying Data Validity, Evaluation Validity and Predictive Modeling for Single-cell Perturbation Simulation

**DOI:** 10.64898/2026.10.13.758713

**Authors:** Tianyi Xia, Lei Cao, Nanning Chen, Haiyang Hou, Changhao Gong, Huan Zhang, Hua Qin, Luni Hu, Di Wang, Qingsong Li, Yong Zhang, Shuangsang Fang, Yuxiang Li

## Abstract

Single-cell perturbation profiling is limited by cost and throughput, while computational methods remain constrained by inconsistent evaluation and underperformance relative to simple baselines. Here we present DCAF, a unified framework for single-cell perturbation simulation that jointly realizes data validity, evaluation validity and modeling predictability. Among its three components, DCAF-data addresses the single-measurement limitation by reconstructing pre- and post-perturbation status using entropy-regularized optimal transport, and is demonstrated on both cell line and multi-cell type datasets; DCAF-criterion evaluates log fold changes in non-silenced genes to improve sensitivity to model error and robustness to zero inflation noise, validated on synthetic datasets; while DCAF-model combines JEPA with explicit modeling for regulatory-dynamics modeling for interpretable prediction. Across benchmarks, our method outperforms state-of-the-art deep-learning methods, as well as mean and linear baselines in cross-gene, cross-cell-type and cross-region generalization. In adult Mouse brain, it recapitulates known cell-type-specific regulatory heterogeneity of Dgcr8 knockdown at both cell type and gene levels. DCAF thus provides a rigorous and biologically grounded foundation for reliable perturbation simulation across development, disease and therapeutic discovery.

## Introduction

One of the focuses of current single-cell functional genomics is to understand how state and function of a cell change in response to perturbations[1][2]. Biotechnologies such as multiplexed Clustered Regularly Interspaced Short Palindromic Repeats (CRISPR)[3] and Perturb-seq[4] provide us with insights into gene regulations in such scenarios yet leading to gigantic gene-cell type-context spaces to be measured. Hence, *in silico* prediction methods are urgently needed and being developed, to serve as a computational complement by enabling simulating perturbation of different unseen genes, cell types, or even regions, guiding cost-effective experiment designs[5][6][7]. Three directions have been developed to categorize most prediction methods[8][9][10], including mechanism-driven methods adopting predefined kinetic mechanisms[11][12][13], constraint-driven methods using restrictions to characterize cells’ behavior ranges[14][15][16], and data-driven paths fitting complex regulation processes through large amounts of parameters and training cells[17][18][19][20].

However, three problems are present to largely affect development of this field. First, as a single cell can only be measured once by current technologies, regulation patterns learned by current models largely involve heterogeneity between randomly matched control and case cells [21][22]. Secondly, diverse measurement criteria capture distinct aspects of method accuracy, and often lead to divergent conclusions. For instance, count-based metrics like MAE[23] measure overall predicted expression profiles, reference-based metrics such as Pearson of change[17][19] and Systema[24] calibrate before measuring predicted expression by referring to control or central baselines, while DEG-based metrics[17] imposes spotlight on biological signals and emphasize if method can specify those signals or not (Supplementary Fig. 1). Zero inflation part of normalized transcriptomics introduces non-biological noise, that harms assessment sensitivity and specificity by forcing additional consideration on genes with subtle changes yet do not affect prediction of cell’s activities. Yet no measurement has been taken to avoid such influences. Thirdly, across all measurements, systematic benchmarks show that current methods do not consistently outperform simple linear or mean baselines, even for complex deep-learning models and foundation models[24][25]. This indicates current methods do not necessarily capture processes describing a cell’s regulation. Some existing methods claim to mitigate this problem by adopting Maximum Mean Discrepancy (MMD)[26] as loss function, yet without adequate reasoning or proof[19], and minimizing MMD in mini-batch training does not lead to single cell accuracy, as a batch of random control and case cells may be drawn from different distributions.

**Figure 1.**
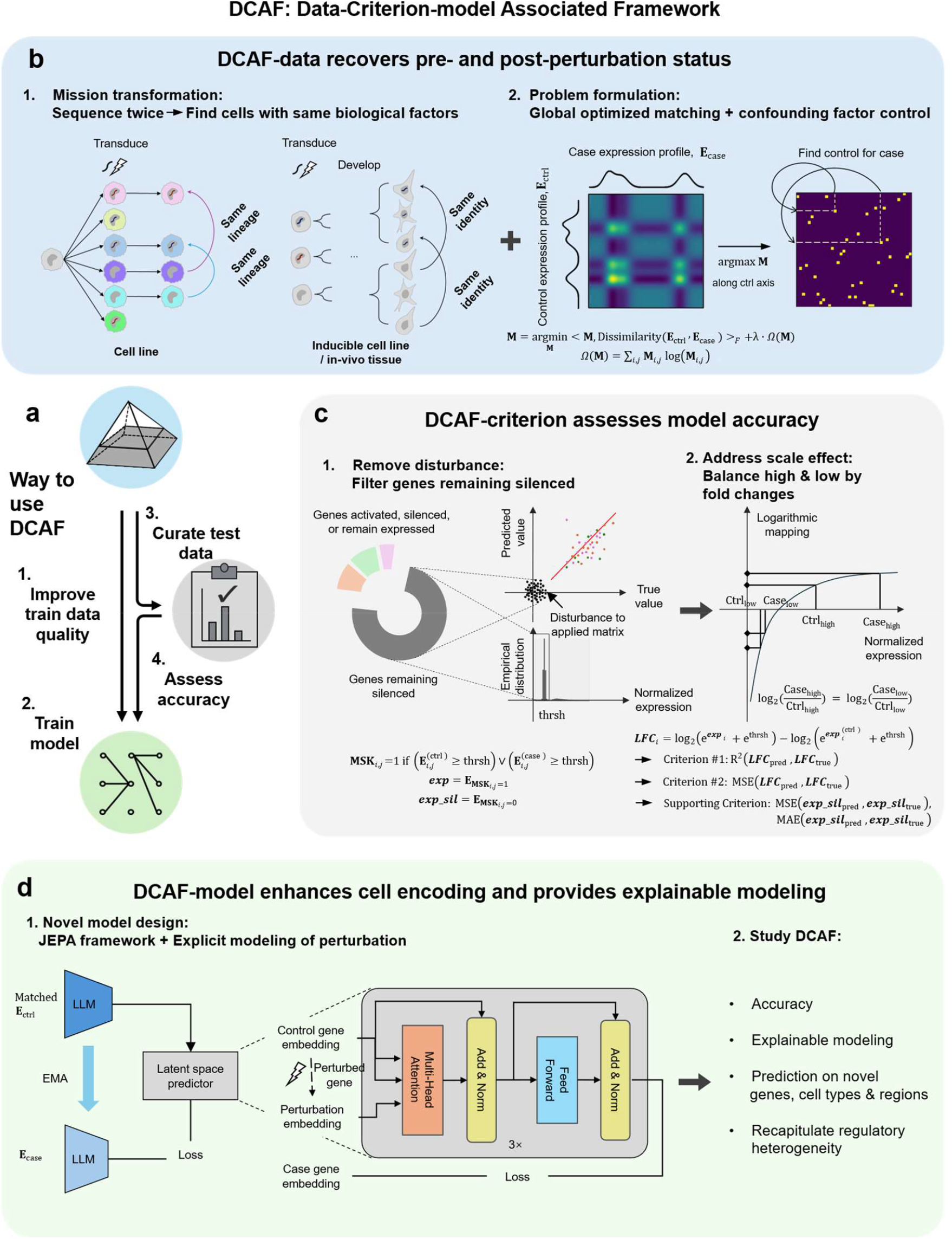
Overview of the Data-Criterion Associated Framework (DCAF) for single-cell perturbation prediction model development. Surrounded by dashed lines are components of DCAF design. (a) DCAF’s components and workflow to use DCAF. (b) DCAF-data that recovers paired pre- and post-perturbation cell states. DCAF leverages shared lineage and cell identity across biological samples to match a control cell with same biological factors for each case cell. It adopts entropy-regularized optimal transport to optimize cell matching and constrain redundant one-to-many pairing. (c) DCAF-criterion raises evaluation standard across divergent assessment standards. DCAF-criterion uses empirical thresholding to mask persistently silent genes and exclude zero-inflation interference. Log fold-change transformation corrects expression magnitude bias across genes with divergent abundance. Genes that remain silent and those that do not remain silent are measured separately by DCAF-criterion. (d) DCAF-model based on JEPA. This predictor adopts Joint Embedding Predictive Architecture (JEPA) and explicit perturbation modeling. DCAF-model was studied through cycle of its training, benchmarking and application.

To address these challenges, we propose the Data-Criterion-model Associated Framework (DCAF), operating at three levels: DCAF-data, DCAF-criterion, and DCAF-model. At the data level, DCAF-data reframes the task of recovering control-case status as finding, for each perturbed cell, an unperturbed cell with near-consistent biological factors. Control and case cells are matched via entropy-regularized optimal transport based on transcriptomic similarity, while accounting for confounding factors including cell composition and batch effects. At the measurement level, DCAF-criterion measures R^2^ and MSE of log fold change between true and predicted expression, focusing on genes that do not remain silenced to prevent uninformative genes from dominating accuracy assessment, and a separate MAE and MSE metric is used for remain-silenced genes. This establishes a perturbation-sensitive, noise-robust criterion for developing, assessing, and selecting perturbation prediction models. At the model level, DCAF-model adopts a Joint Embedding Predictive Architecture (JEPA) to enhance the encoder’s understanding of pre- and post-perturbation status and incorporates the first neural network that explicitly models regulatory dynamics, providing interpretable and reliable prediction at the full transcriptome level.

We validate DCAF both analytically through studying each of its three components, and experimentally through applying DCAF-model on Human brain organoid and adult Mouse brain perturbations. For DCAF-data, accurate recovery of pre- and post-perturbation status recovery was witnessed on 3 multi-cell types and 2 cell line data. When studying efficacy of DCAF-criterion, a far higher sensitivity to error, and specificity to disturbance was found, on synthetic data carrying known levels of error. DCAF-model was validated through effects of its JEPA structure and explicit modeling of regulation process. Then its accuracy was demonstrated using 5 cell-line and 3 multi-cell type datasets, with an obvious advantage compared to autoencoders, LLM and simple linear and mean baselines. On Human brain organoids, model directed by DCAF accurately predicts both novel genes and novel cell types. When generalizing on a novel MOp region of adult Mouse brain, the trained model is found to recover both cell-level and gene-level regulatory heterogeneity consistent with known neurobiological mechanisms. To conclude, DCAF provides a rigorous foundation spanning data quality control, model development, and assessment, thereby enabling the reliable computational predictions essential for target prioritization, disease modeling, and experiment design[4][27][28].

## Results

### Design of DCAF

DCAF is a designed foundation to predict perturbation results on single cells, from data, criterion and model perspectives. DCAF-data improves model performance through enhancing quality of training data and enhances model assessment reliability by curating the testing dataset (Fig.1a). By DCAF-criterion, model accuracies are sensitively evaluated, with guarded robustness in presence of noise disturbance. Through DCAF-model, exceeding accuracy is reached, enabling prediction crossing gene, cell types and regions.

### DCAF-data for data preparation for training and assessment

In DCAF-data framework, we leverage Entropy-regularized Optimal Transport (OT)[29][30] to recover pre- and post-perturbation single cell states formalized as a connectivity map (Methods). DCAF-data transforms the almost impossible task of finding control status, to finding an unperturbed cell whose other biological factors are almost consistent with a perturbed cell (Fig. 1b). Its global setting suppresses one-to-many pairings due to confounding factors[22] by optimizing overall pairwise transcriptional similarity. And entropy regularization avoids forced employment of all control cells. After solving the connectivity map, for each case cell, the control cell with highest probability is picked up as its matched control status. Critically, this matching process is applied both on cells fed to training model, and on cells used in testing phase, establishing a standardized, biologically validated data foundation for both simulator development and measurement (Fig. 1a).

### DCAF-criterion for prediction accuracy assessment

When assessing prediction accuracy of methods, DCAF-criterion first implements thresholding strategy based on empirical distribution of gene expression of the specific dataset, to avoid disturbances of genes that remain silent throughout perturbation for each cell (Fig. 1c, and Methods). The thresholding operation eliminates interference of those genes on performance metrices, thereby significantly improving the above-mentioned sensitivity and specificity of model evaluation. Next, DCAF-criterion counts log fold change of all non-silenced transcripts for each cell, to balance influence of varying expression levels of each gene and emphasize the prediction of regulation result. Both R^2^ and MSE are used to score the accuracy of model, with R^2^ of log fold change = R^2^(***LFC***_pred_, ***LFC***_true_) and MSE of log fold change = MSE(***LFC***_pred_, ***LFC***_true_). For genes that remain silenced for each cell, a custom MSE and MAE matrix are used to measure model’s performance. The above criteria enhance assessment sensitivity by outlining model’s performance on low-abundance transcripts and improve specificity by equally considering genes with different expression levels.

### DCAF-model for modeling single-cell level regulation in response to perturbation

Unlike current methods which most adopt generative framework where only control cells are projected to their latent space, DCAF-model adopts and re-adjust the joint embedding predicative (JEPA)[31][32] framework, where control and case cells respectively go through embedding models with almost identical parameters controlled by Estimated Moving Average (EMA) [32], and a predictor is adopted to predict control gene embedding along with perturbation information, to the projected case gene representation (Fig. 1d, Supplementary Fig. 2 and Methods). Such framework allows encoder to receive both pre- and post-perturbation as inputs, hence largely increases its understanding of the perturbation dataset. Pretrained scGPT[18] is adopted as the encoder, with Low-Rank Adaptations (LoRA)[33] applied for its finetuning. The predictor model is essentially a 3-layer transformer model with the first block implementing cross-attention between perturbation information and control gene embeddings, and last two blocks applying self-attention of projected gene embeddings. The cross attention encodes stimulation of gene perturbation as product of Q and K measures correlations across highly variable genes, while perturbation information encoded in V is extracted by the product to assign its initial influence onto gene regulatory networks. The self-attention blocks mimic the self-regulation process of each perturbed cell, leading to a final perturbed embedding. Reconstructor training is implemented in the following phase, where the encoder and predictor are frozen, and only a reconstructor is trained to project control cells along with perturbation information to their case gene expression. Training data of DCAF-model is preprocessed by DCAF-data to recover pre- and post-perturbation status.

With DCAF-data, DCAF-criterion and DCAF-model combined, a solid foundation is provided for perturbation prediction, with basis data quality ensured, method selection standard established, and model reliability being increased.

### DCAF provides data and criteria basis for perturbation modeling

As DCAF-data provides data basis to prepare both train and test datasets for models, and DCAF-criterion provides assessment standards to benchmark trained models and guide model selection, their effects as supporting foundation are studied for perturbation modeling (Fig. 2a).

To evaluate DCAF-data, we benchmarked it against random matching, random matching within each cell type, and CINEMA-OT[22] on both multi-cell-type datasets (*in utero*[34] and adult Mouse brain[35], Human brain organoid[36]) and neuronal cell lines with 1–2[37] and 1–5 genes perturbed per cell[37]. All datasets derive from healthy cells under the assumption that perturbation leaves most of their genes unaffected. On multi-cell-type data, DCAF achieves the lowest mean KL divergence (Fig. 2b); on cell lines, it yields markedly reduced divergence relative to baselines. Ablation revealed that removing global optimization causes the proportion of matched control cells to drop below unrealistic 20% in four of seven datasets (Fig. 2c, Supplementary Fig. 3, Supplementary Table 1). Meanwhile, removing the regularization term drives the matched control ratio to nearly 1.0 in five of seven datasets, contradicting the expectation that not all control cells have a perturbed counterpart in a given experiment. To assess the influence of DCAF-data on pre- and post-perturbation status recovery, we examined its performance on both cell line and multi-cell-type data. On neuronal cell lines with 1 to 2 and 1 to 5 genes perturbed per cell , DCAF-matched cell pairs exhibit a KL divergence distribution closer to zero relative to random matching (Fig. 2d), indicating that cells with higher expression similarity are paired between pre- and post-perturbation states. On the *in utero* Mouse brain dataset, control cells clustered by their correlation with each case cell recapitulate the clustering structure of case cells, demonstrating that hierarchical cellular organization is preserved through DCAF-data recovery (Fig. 2e). Marker gene enrichment analysis further reveals strong concordance between DCAF-matched control cells and their corresponding case cell types, mirroring the enrichment pattern of the case cells themselves, whereas no such correspondence is observed under random matching. Consistent results were also obtained on the Human brain organoid dataset (Supplementary Fig. 4), confirming that both hierarchical structure and gene enrichment patterns are faithfully recovered by DCAF-data when reconstructing pre- and post-perturbation status.

**Figure 2.**
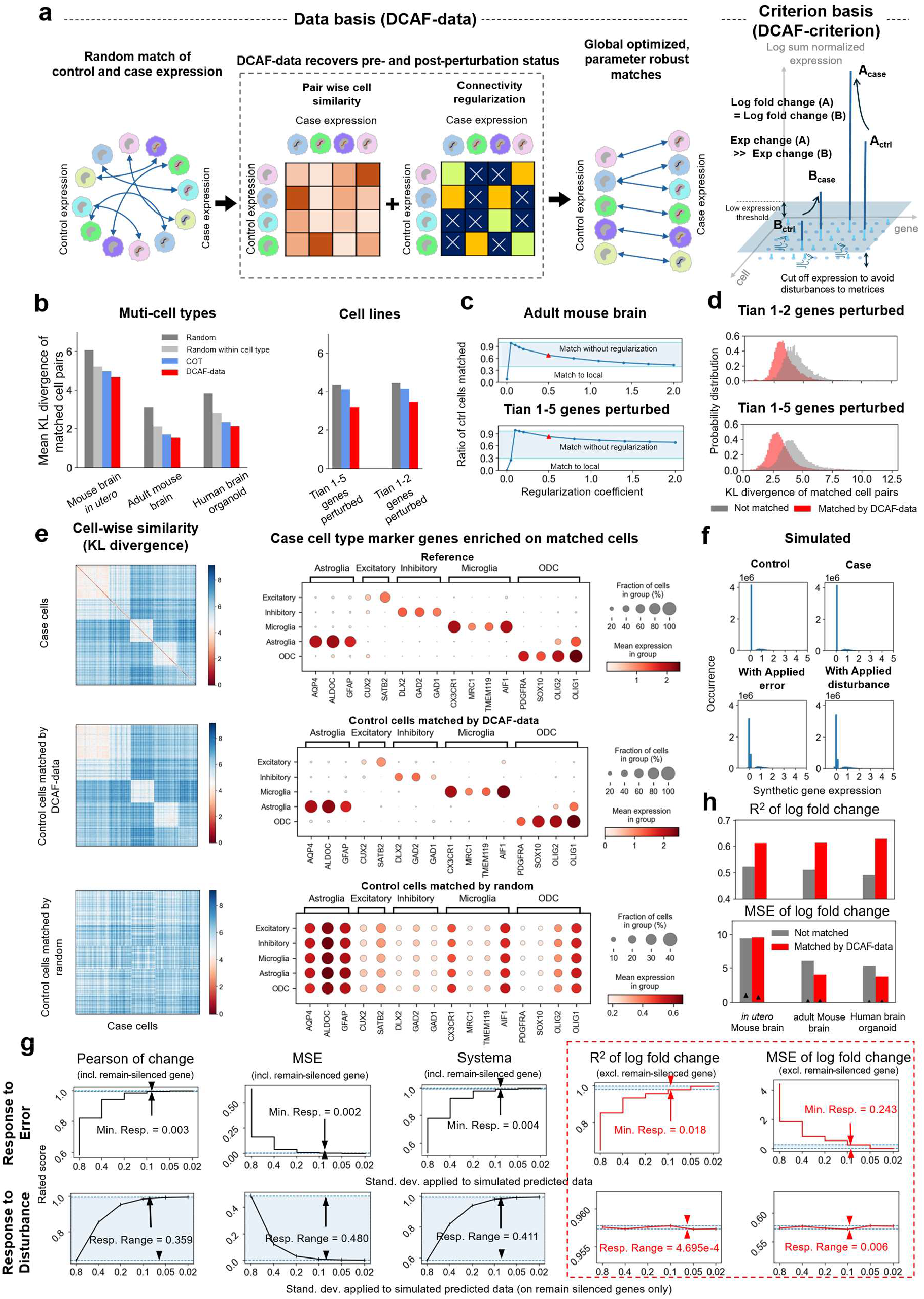
DCAF provides data and criteria basis for perturbation modeling. (a) DCAF-data’s stepwise cell pairing pipeline from raw expression toward globally optimized case-control matching, and DCAF-criterion’s measuring process removing noise from persistently silenced genes and balancing importance of different expression levels. Surrounded by black dashed lines are DCAF’s recovery steps. (b) Comparison on recovery accuracy by different methods across multi-type brain tissues and perturbed cell-line datasets. (c) Ratio of matched control cells caused by matching without regularization, local matching, and matching by DCAF. The upper arrow marks matching without regularization and lower arrow denotes locally restricted matching. Red triangle highlights ratio of matched control cells caused by fixed regularization parameter by DCAF. (d) Probability distribution of pairwise KL divergence calculated from matched case-control cell pairs under DCAF matching and random matching. (e) Comparison of intrinsic cell hierarchy and marker gene enrichment on *in utero* Mouse brain recovered by different methods. Present from left to right are KL divergence heatmaps and marker enrichment plots. From top to bottom presents recovered information from different sources, including case cells themselves, control cells matched by DCAF, and randomly matched control cells. (f) Expression distributions of selected synthetic data. Presented on the 1^st^ row are simulated control and case expression distribution. Shown in the 2^nd^ row are simulated prediction datasets with known levels of error and disturbance, with standard deviation of 0.05 in Gaussian noises. (g) Comparative responses of multiple evaluation metrics toward gradually refined disturbance gradients, including Pearson of change, MSE, Systema, R^2^ of log fold change, and MSE of log fold change. Surrounded by red dashed lines are DCAF’s matrices. (h) Cross-dataset performance comparison between our model architecture trained on DCAF-matched and randomly matched cell pairs. Plotted as bars are measurements of prediction on genes not remaining silent. Plotted as black triangles are measured MSE of prediction on genes that remain silent.

To evaluate DCAF-criterion, we assessed its sensitivity to model error and robustness to disturbances using simulated datasets with controlled error levels (Methods). Control cells were generated from a zero-inflated binomial distribution calibrated to real datasets, and case cells were synthesized with fraction of activated, silenced, and remaining-expressed genes consistent with real-world datasets [38][39][35][27][40][36] (Supplementary Fig. 5, Supplementary Fig. 6). Gaussian noise of exponentially varying standard deviation was applied to simulate model outputs at different error levels (Fig. 2f, Supplementary Fig. 7). DCAF-criterion, which measures R^2^ and MSE of log fold change excluding remain-silenced genes, was compared against Pearson correlation of expression change, MSE, and Systema. DCAF-criterion exhibited minimum responses 3.5-fold and 60-fold higher than Systema, which is the best-performing existing method, on R^2^ and MSE of log fold change respectively (Fig. 2g), demonstrating markedly higher sensitivity. This advantage stems from log fold change-based evaluation, which equally weights genes across expression levels, whereas expression-based metrics disproportionately emphasize highly expressed genes (Supplementary Fig. 8). Notably, applying remain-silenced gene exclusion to existing metrics did not improve their sensitivity (Supplementary Fig. 9), confirming that DCAF’s advantage derives from equal treatment of genes at different expression levels. For genes remaining silenced, similar results were shown by its response (Supplementary Fig. 10). For robustness, disturbances were simulated by injecting Gaussian noise into genes remaining not expressed only. DCAF achieved the most stable scores across disturbance levels, with response ranges 898-fold and 60-fold lower than Pearson correlation for R^2^ and MSE of log fold change, respectively. This robustness is attributable to the exclusion of remain-silenced genes, which filters out outlier log fold changes from extremely-low-expression genes that disproportionately affect evaluation without reflecting genuine model performance. Reintroducing remain-silenced genes sharply increased the response range of DCAF metrics, and applying this exclusion to existing methods correspondingly improved their robustness (Supplementary Fig. 11). Applying DCAF-criterion to models trained with DCAF-matched versus randomly matched data revealed a clear improvement in accuracy on non-silenced genes across *in utero* and adult Mouse brain and Human brain organoid datasets (Fig. 2h), while scores on remain-silenced genes remained nearly constant, validating the effectiveness of DCAF-data in improving model performance.

### DCAF-model achieves advanced accuracy over existing approaches on cross-gene and cross-cell-type perturbation prediction

To study effectiveness of innovative components of DCAF-model, we first compare DCAF-model’s JEPA architecture with custom generative framework under identical conditions (Methods). On H1 HESCs, where perturbations exert large effects on gene expression[40], cells with different target genes occupy more separable regions by DCAF-model than generative framework (Fig. 3b). While more integrated per-cell-type manifolds are observed on both adult and *in utero* Mouse brain datasets by DCAF-model than model with generative framework (Fig. 3c, Supplementary Fig. 12). Quantitatively, DCAF-criterion of genes not remaining silenced reveals markedly improved accuracy under DCAF-model on Frangieh[38], H1 HESCs[40], and adult Mouse brain[35] (Fig. 3d). Meanwhile, MSE on remain-silenced genes remains consistent between frameworks. Above findings demonstrate DCAF-model’s advantage in capturing cell context by adopting JEPA structure, as its encoder is trained to encode both control and case transcriptomics.

**Figure 3.**
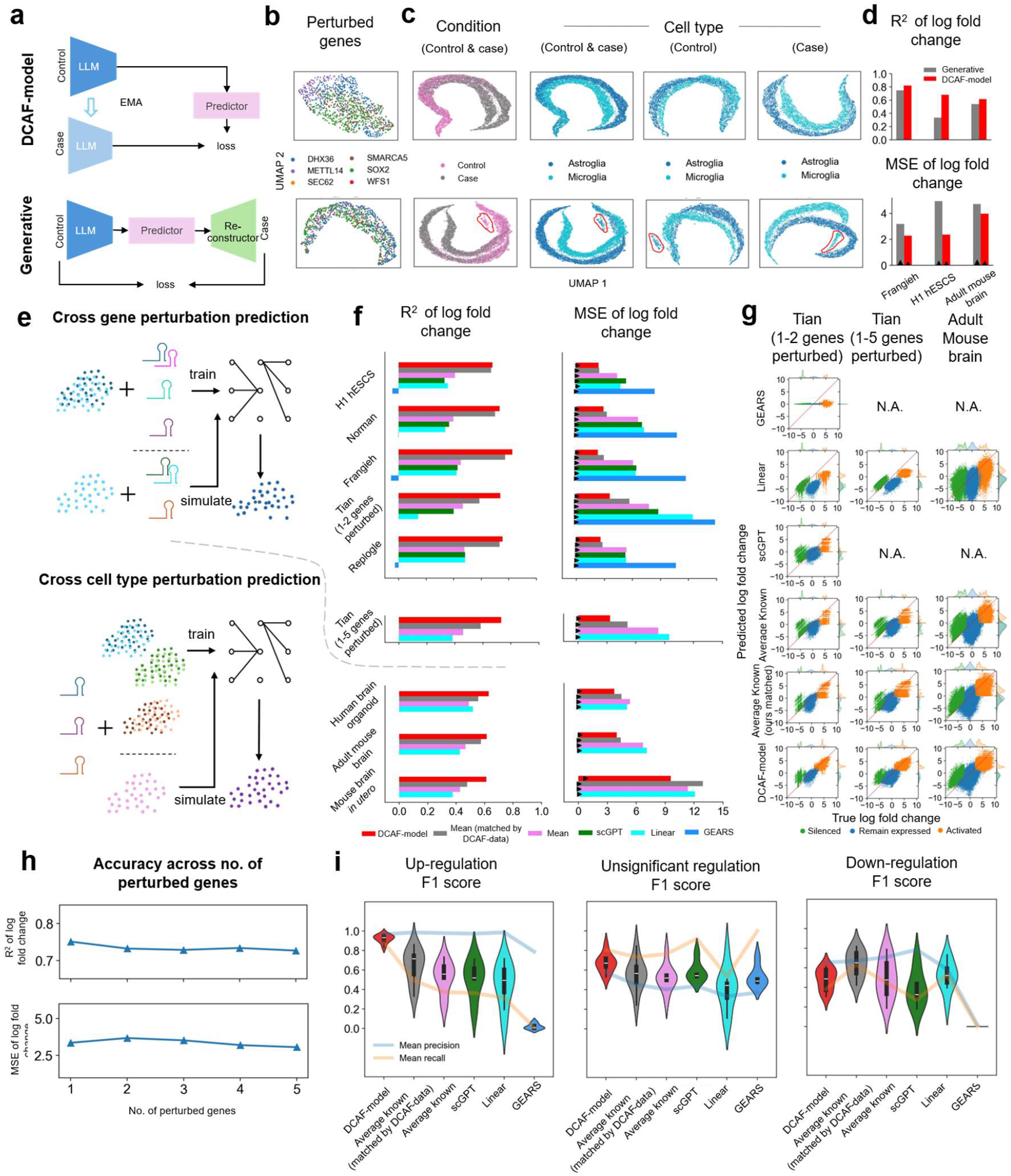
DCAF-guided model is validated, then benchmarked against existing approaches across cross-gene and cross-cell-type prediction scenarios, using both DCAF criterion and conventional non-indexed metrics. (a) Comparison between DCAF-model framework and generative framework. Presented in top rows of (a), (b) and (c) are DCAF-model structure and results, while in bottom rows are generative framework structure and results. (b) UMAP manifold of H1 hESCS test cells encoded by DCAF-model and generative framework. (c) Manifold of *in utero* Mouse brain test cells encoded by DCAF-model and generative framework. Shown in the 1^st^ and 2^nd^ columns are tested control and case cells colored by condition and cell types. Shown in the 3^rd^ and 4^th^ columns are respectively control and case cells colored by cell types. (d) DCAF-accuracy of DCAF-model and generative model on Frangieh, H1 hESCS and adult Mouse brain datasets. Plotted as bars are measurements of prediction on genes not remaining silent. Plotted as black triangles are measured MSE of prediction on genes that remain silent. (e) Definition of predicting scenarios including cross gene perturbing prediction and cross cell type perturbing prediction. (f) DCAF-criterion assessment of methods including GEARS, linear baseline, scGPT, mean baseline and DCAF-model, across different cell lines and multi-cell types datasets. (g) Comparison between predicted and actual log fold change of silenced, persistently expressed and activated genes for all compared methods in (f). (h) Changes in the model’s DCAF-criterion scores alongside varying numbers of perturbed target genes. (i) F1 score distributions, mean recall and mean precision for up-regulated, non-significantly regulated and down-regulated genes, across all datasets, and by all approaches in (f).

We then examine DCAF-model’s explainability through its innovative explicit modelling predictor. Attention maps across transformer layers were visualized on Frangieh, Tian[37], and H1 HESCs datasets. Gene clusters exhibit more uniform attention values in the third layer versus the second (Supplementary Fig. 13), indicating clearer gene cluster discrimination from lower to upper layers. Hierarchical gene clusters from the third attention layer yield apoptotic, neuronal, and morphogenesis enrichment for the three datasets, respectively (Supplementary Fig. 14), suggesting progressive feature aggregation within gene modules along model depth. Min-max scaled and Gaussian-denoised attention distributions show a wider range and lower peak from the second to third layer (Supplementary Fig. 15), indicating increasingly diverse attention patterns across genes at deeper layers.

With innovative components validated, we benchmark accuracy of DCAF-model through comparing with other current methods. After recovering pre- and post-perturbation status via DCAF-data, matched training data were used to train our perturbation prediction model, and DCAF-criterion was applied to compare performance between DCAF-model and existing deep learning approaches on matched test data. Two scenarios were evaluated: cross-gene perturbation simulation and cross-cell-type perturbation simulation. In the cross-gene scenario, perturbations on a limited set of genes are seen during training, and unseen perturbations are used to test generalization on the same cell line (Fig. 3e). Both datasets with 1–2 target genes[40][39][38][37][27] and one dataset with 1–5 target genes[37] are included. In the cross-cell-type scenario[36][35][34], perturbations on a subset of cell types are used for training, while the trained model is tested on unseen cell types with the same perturbed genes. Measured by DCAF-criterion on genes not remaining silent, DCAF-model achieves the highest accuracy on both R^2^ of log fold change and MSE of log fold change, compared to GEARS, scGPT, linear baseline, and the mean baseline, which uses the mean perturbation result of training data as predicted transcriptomics (Fig. 3f). MSE on remain-silenced genes is largely consistent across methods. The proximity between DCAF’s MAE measurement and the low threshold of these genes, which share the same dimension, indicates that DCAF-model’s prediction on remain-silenced genes reaches advanced accuracy (Supplementary Table 2). An optimal accuracy by DCAF-model is also observed across silenced genes, activated genes, and genes that remain expressed (Fig. 3g). Measured from a gene regulation perspective, the same accuracy ordering among methods is observed relative to DCAF-criterion, on both up-regulation and non-significant regulation F1 scores (Fig. 3i, Methods), reconfirming the reliability of DCAF-criterion in methods evaluation. Comparable results are achieved across methods on down-regulation prediction. When training data are preprocessed by DCAF-data, DCAF-model still surpasses mean baseline, the best-performing approach, by both DCAF-criterion and regulation measurements (Fig. 3f, Fig. 3i, Methods), confirming that the accuracy increasement stems from model design rather than training data quality. A steady score is observed when measuring DCAF-model’s performance across varying numbers of perturbed genes, indicating prediction stability (Fig. 3h).

### DCAF-model predicts perturbation outcomes for both novel genes and novel cell types on Human brain organoid

To evaluate the generalization capability of DCAF-model, we designed an intra-tissue transfer prediction task involving prediction under novel perturbed genes and on novel cell types. Specifically, using the Human brain organoid dataset[36], the model was trained on a subset of perturbed genes and tested on unseen gene perturbations, as well as trained on specific cell types and tested on unseen cell types (Fig. 4a).

**Figure 4.**
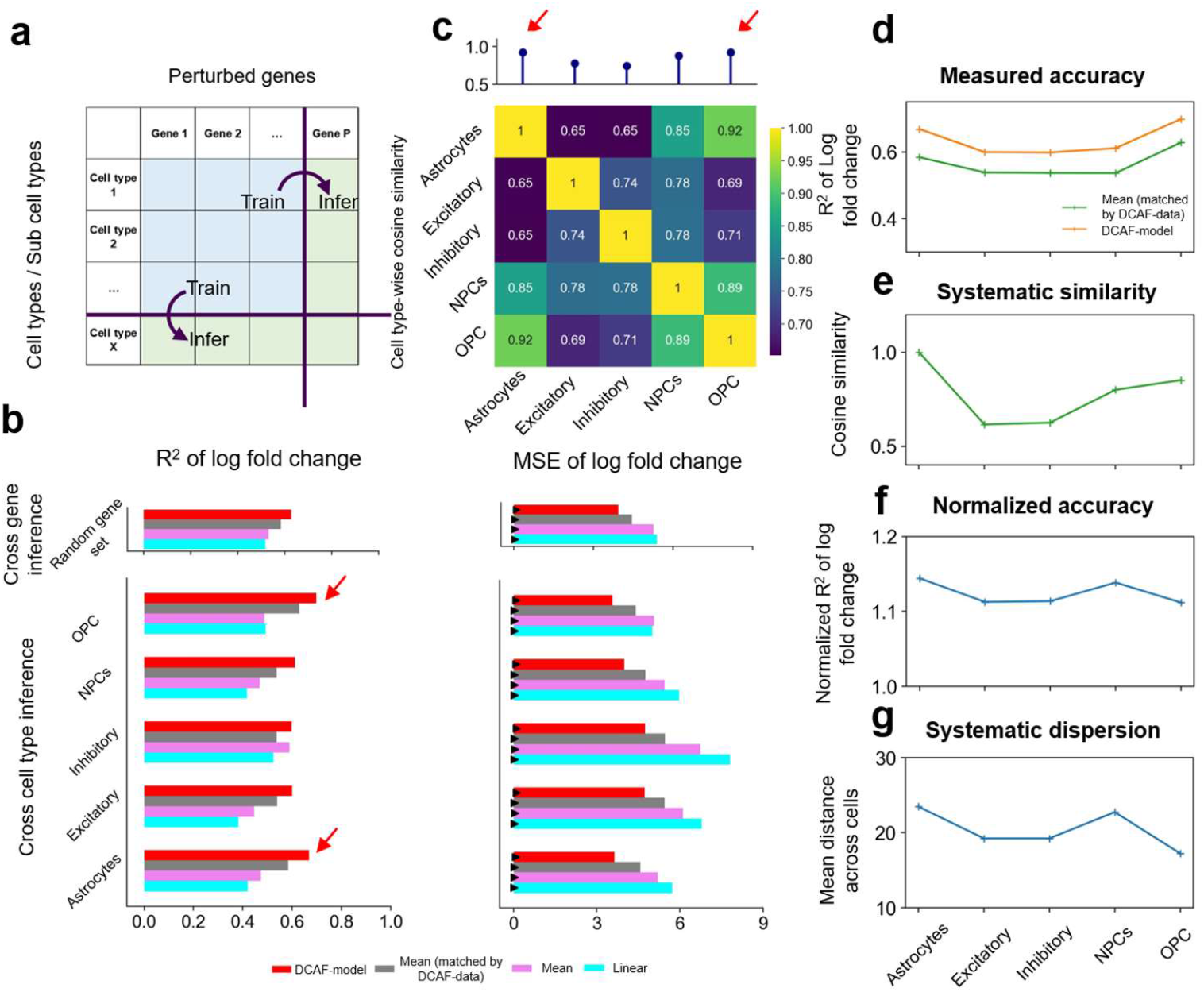
DCAF-guided model realizes effective generalization to unseen perturbed genes and unseen cell types on Human brain organoid dataset. (a) Experimental design of two generalization prediction tasks: cross-gene inference and cross-cell-type inference split for training and testing. (b) DCAF accuracy comparison between linear baseline, mean baseline and DCAF-guided model, under cross-gene and cross-cell-type prediction conditions. Plotted as bars are measurements of prediction on genes not remaining silent. Plotted as black triangles are measured MSE of prediction on genes that remain silent. (c) Cosine similarity between cell types, with maximum cross-cell type similarity value shown above. Red arrows mark the highest two maximum similarity values. (d) Measured DCAF R^2^ accuracy of DCAF-guided model and mean baseline across distinct held-out cell-type groups. (e) Systematic similarity values corresponding to each tested cell type. (f) Normalized accuracy of DCAF-guided model against mean baseline, across distinct held-out cell-type groups. (g) Systematic dispersion values corresponding to each tested cell type.

Under these settings, DCAF-model achieves the best performance across both tasks (Fig. 4b). For unseen gene prediction, on genes not remaining silent, the model attains the highest R^2^ and the lowest MSE of log fold change across randomly sampled gene sets, outperforming linear and mean baselines both with and without DCAF-matched training data. On remain-silenced genes, MSE is largely consistent across methods. For the held-out gene KDM6A, predicted and true log fold change distributions are closely aligned (Supplementary Fig. 16). Above results indicate that DCAF-model captures transferable perturbation patterns beyond the observed gene space rather than relying on interpolation or averaging. For unseen cell type prediction, on genes not remaining silent, the model consistently achieves optimal performance across all leave-out cell types, including Astrocytes, Excitatory neurons, Inhibitory neurons, NPCs, and OPCs, while error on remain-silenced genes remains largely constant across methods and cell types. These findings demonstrate that DCAF-model generalizes perturbation response patterns from observed to previously unseen cell types. To investigate the basis of this performance, we examined the relationship between dataset similarity and prediction accuracy across train-test splits. A cell-type similarity matrix reveals correlations between tested and training cell types (Fig. 4c, Supplementary Fig. 17). Astrocytes and OPCs exhibit higher maximum cosine similarity to single trained cell types relative to other splits (Methods), consistent with the overall prediction advantage in the cross-cell-type task, suggesting that the model preferentially transfers learned perturbation patterns to target cell types with similar transcriptional profiles.

To investigate the mechanisms underlying this advantage, we adopt Mean baseline[25] as the reference baseline and provided both DCAF-model and reference baseline with cells matched by DCAF-data. To adapt Systematic Variation framework from Systema[24] to the train-test-split level, we measured the distance between centroid of aggregated trained cell types and each leave-out test type (Methods). When Astrocytes and OPCs are left out as test types, higher systematic similarity and higher accuracy co-occur across methods (Fig. 4d, e). This is consistent with Systema’s finding that higher systematic variation yields improved prediction. Normalizing DCAF scores over the baseline, the highest normalized accuracies occur when Astrocytes or NPCs are left out (Fig. 4f). Systematic dispersion of test cell types mirrors this trend (Fig. 4g), suggesting that greater transcriptomic dispersion confers a larger advantage to DCAF-model. This indicates cellular heterogeneity is captured by the model to enable single-cell-level prediction accuracy. Collectively, above results demonstrate that DCAF-model transfers perturbation predictions across genes and cell types at single-cell resolution.

### DCAF-model predicts perturbation outcomes in novel brain regions

To evaluate the generalization capability of DCAF-model across distinct brain regions, we trained on Mouse prefrontal cortex (PFC)[35] and transferred to primary motor cortex (MOp)[41], which differ in both function and cellular composition (Fig. 5a, Methods).

**Figure 5.**
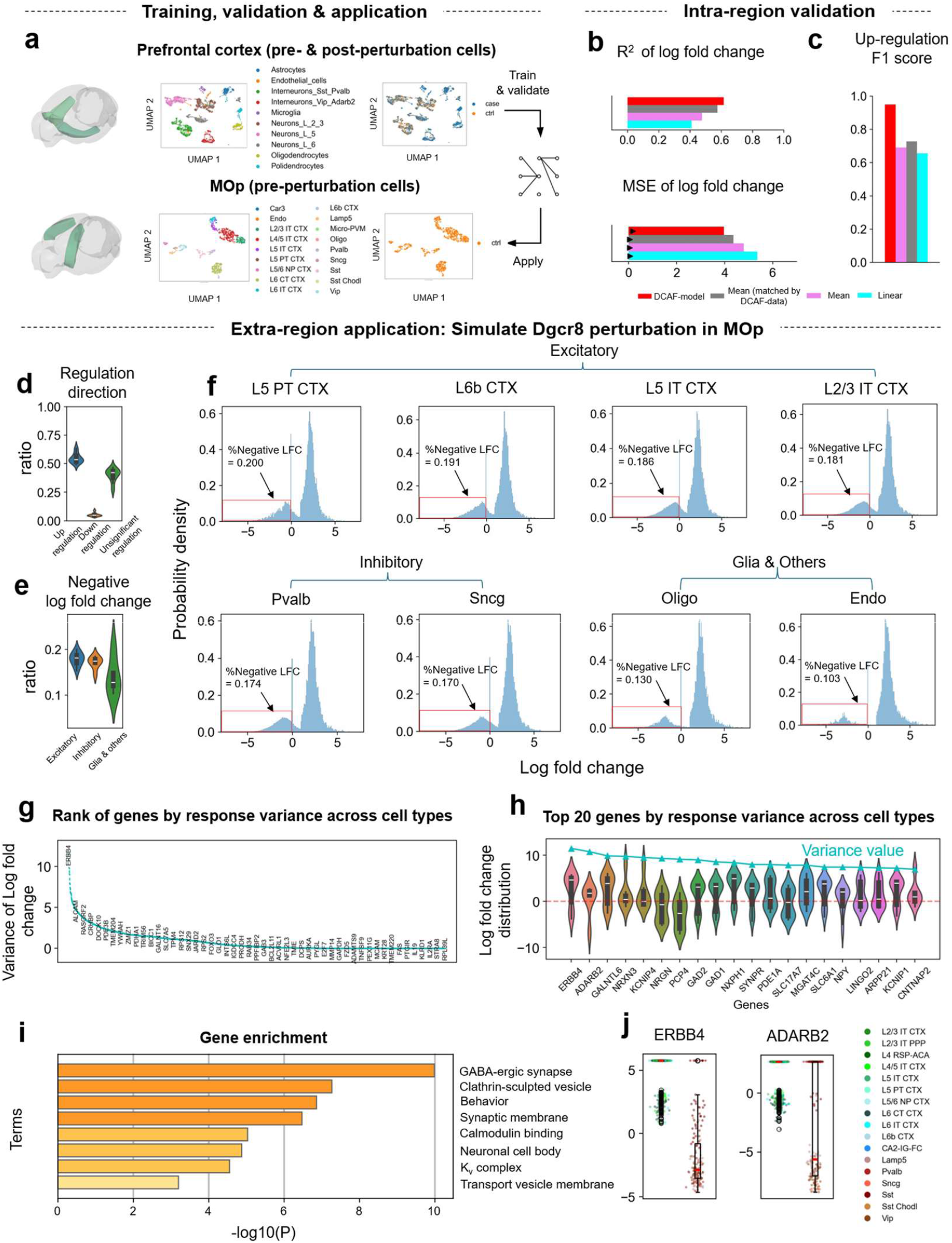
DCAF-guided perturbation prediction model generalizes across brain regions of adult Mouse brain. (a) Overview of the cross-brain-region generalization task. The model is trained and validated on prefrontal cortex (PFC) perturbation dataset and transferred to MOp region. The 1^st^ and 2^nd^ row respectively shows UMAP of prefrontal cortex and MOp cells. The 2^nd^ column and 3^rd^ column are colored respectively to highlight if each cell is perturbed (marked as case) or not (marked as ctrl). Only 300 cells are randomly selected and plotted for each cell type with over 300 cells when plotting the top row, for a clear visualization. (b) DCAF accuracy compared between Linear, Average known and DCAF-guided model, under cross-cell-type prediction conditions. Plotted as bars are measurements of prediction on genes not remaining silent. Plotted as black triangles are measured MSE of prediction on genes that remain silent. (c) Up-regulation classification performance (F1 score) of the DCAF-selected model versus baseline methods including Linear and Average known. (d) Distribution of up-regulation, down-regulation and unsignificant regulation ratio of predicted responses to Dgcr8 perturbation, across different cell types in MOp. The whiskers extend from the box to the farthest data point lying within 1.5 times the inter-quartile range (IQR) from the box. Median values of each data are highlighted by white solid lines. (e) Distribution of ratio of negative predicted log fold change across cell types, for excitatory, inhibitory, and glial/other neurons. Whiskers and white solid lines represent the same meaning as described in (d). (f) Distribution of log fold change across different genes of different cells, for 4 cell types for excitatory neurons, 2 cell types of inhibitory neurons, and 2 cell types for glial/other neurons. Negative log fold change part is squared by red solid lines, pointed to from specific ratio statistics. (g) Ranking of genes by their response variance, measured by variance of mean LFC across different cell types, with 50 representative genes labeled at fixed intervals. (h) Distribution of log fold change values across cell types for the top 20 genes ranked by cross-cell-type response variance. The variance of log fold change is indicated by the blue line, with the specific variance value for each gene marked by blue triangles. Whiskers and white solid lines represent the same meaning as described in (d). (i) Functional enrichment analysis of the 20 genes shown in (h). (j) Single-cell resolution predicted responses for the top two high-variance genes, Erbb4 and Adarb2, showing divergent regulation in excitatory versus inhibitory neurons. Whiskers represent the same meaning as described in (d). Median values of each data are highlighted by red solid lines.

Internal validation within PFC, which holds out a subset of cell types as the test set, confirmed robust generalization. Specifically, compared to baselines, DCAF-model achieves significantly higher R^2^ and lower MSE of log fold change on genes not remaining silent, along with the highest F1 score on both upregulated and insignificantly regulated gene classification (Fig. 5b, c; Supplementary Fig. 18). Error on remain-silenced genes remained consistent across methods.

Building on this validation, we selected Dgcr8 as the target gene for cross-region transfer. Dgcr8 encodes a core component of the miRNA maturation machinery [35]; with haploinsufficiency implicated in 22q11.2 deletion syndrome and causes widespread transcriptional dysregulation[42][43]. At the cell type level, as predicted, Dgcr8 downregulation elicited a bidirectional transcriptional response dominated by upregulation with a subset of downregulated genes (Fig. 5d), consistent with the established mechanism whereby impaired miRNA biogenesis relieves miRNA-mediated repression, leading to global transcriptional de-repression[44]. Across cell types, the proportion of downregulated genes was higher in excitatory neurons, while lower in inhibitory, glia and other neurons (Fig. 5e, f, Supplementary Fig. 19). This gradient aligns with prior evidence that excitatory neurons, with their more complex synaptic architecture and distinct miRNA dependencies, are preferentially sensitive to Dgcr8 loss[45][46].

At the gene level, ranking by cross-cell-type variance in log fold change (Fig. 5g) revealed that many top-ranked genes exhibit opposite regulatory directions across cell types (Fig. 5h). At single-cell resolution, the top two variance-ranked genes, Erbb4 and Adarb2, were predicted to be regulated in opposite directions in excitatory versus inhibitory neurons (Fig. 5j).These opposing responses are biologically plausible given the distinct transcriptional homeostasis and miRNA expression profiles maintained by excitatory and inhibitory neurons[47], which cause the same downstream targets to respond divergently depending on cellular context[46][48]. Functional enrichment of the top 20 genes by response variance includes synaptic transmission, neural signaling, and voltage-gated ion channel processes (Fig. 5i), which are pathways differentially utilized across neuronal subtypes[49].

Collectively, through the Dgcr8 case study, DCAF-model demonstrates the capacity to learn and transfer perturbation prediction to novel tissues at both cell-type and gene levels, likely attributable to the LLM encoder injecting pretrained knowledge of diverse tissues, DCAF-data improving data quality, and DCAF-criterion guiding method selection.

## Discussion

In this work, we establish the Data-Criterion-model Associated Framework (DCAF), a unified, biologically grounded framework to standardize the full lifecycle of single-cell perturbation prediction modeling, spanning data curation, rigorous assessment, and explainable, accurate modeling. The respective components are termed DCAF-data, DCAF-criterion, and DCAF-model. By reconstructing high-fidelity control–perturbed cell pairs via entropy-regularized optimal transport, DCAF-data resolves the fundamental limitation that a single cell cannot be measured before and after perturbation while accounting for confounding effects such as cell composition and batch variation. This capability is demonstrated on both cell line and multi-cell-type datasets. By focusing evaluation on log fold changes of non-silenced genes, DCAF-criterion is shown to enhance assessment sensitivity to errors and robustness to technical noise, outperforming conventional metrics. DCAF-model is constructed using JEPA with scGPT as encoder, which explicitly models regulatory dynamics and achieves superior generalization across unseen genes, cell types, and brain regions, spanning diverse systems including Mouse brain, Human brain organoids, and multiple cell lines. The superiority of DCAF-model is demonstrated by both DCAF evaluation metrics and orthogonal biological perspectives such as regulation direction. The relationship between model performance and systematic similarity and dispersion is examined on the Human brain organoid dataset. Validated in the Mouse brain dataset, the model accurately recapitulates cell-type-specific regulatory heterogeneity across regions, exemplified by Dgcr8 knockdown predictions that align with known neurobiological mechanisms.

DCAF is known to be the first work that reframes single-cell perturbation prediction as a full-stack machine-learning problem, highlighting data validity, evaluation reliability and model interpretability. It emphasizes the essential problems of partially observable data and uninterpretable modelling process, and provides a systematical solution for perturbation modeling development, including data preparation and model assessment, as well as an enhanced model alternative complementing explainability and accuracy. As a result, DCAF establishes a rigorous foundation for reliable and measurable *in silico* perturbation prediction. It paves the way for the development of next-generation data-driven perturbation models that generalize across the vast gene–cell–tissue landscape. Through our study, DCAF is confidently expected to accelerate target prioritization and to support cost-effective experimental design for developmental biology, disease modeling, and therapeutic discovery.

Regarding future work, the improvement in trained model accuracy achieved by DCAF-data underscores the importance of consistent biological factors between input and output transcriptomes during modeling. As being essential for both modelling and analysis, biotechnological approaches, such as sequencing a single cell both before and after perturbation, should be explored to further advance this aspect. An organized utilization of simulation results, towards effective experimental design, can also be explored, as a combination of *in vivo, in vitro* and *in silico* techniques. Furthermore, as higher-volume and more diverse samples are measured and published, a unified perturbation model for a universally studied tissue or cell line, such as K562, is expected to be developed following our framework. Such a model would enable direct inference without the necessity of training, thereby accelerating discoveries in specific biological or medical scenarios.

## Methods

### DCAF-data

According to DCAF framework, single cell transcriptomics of control and case cells are matched, to recover pre- and post-perturbation status by DCAF-data. To start with, we introduce 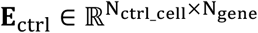 and 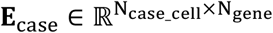 to respectively denote expressions of preprocessed control and case cells. In both matrices, each row represents a cell, and each column denotes a gene. Genes are laid out by the same order in these matrices. The matching of control and case cells is built on the hypothesis that the accumulated similarity among matched cells is higher than any other possible combinations. To formulate this, by DCAF-data, we solve the combination **M** between control and case cells:

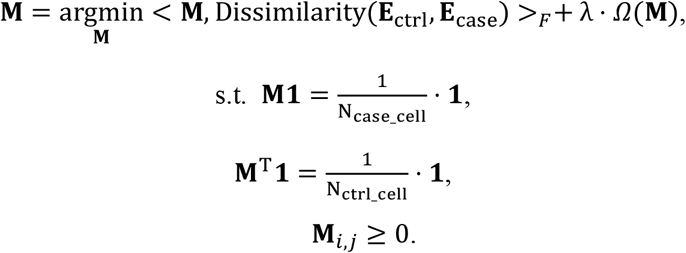

Thereinto,

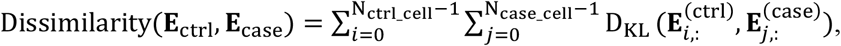

where D_KL_(**X, Y**) denotes the Kullback-Leiber divergence[50] between vectors **X** and **Y**. < **X, Y** >_*F*_ represents the Frobenius product of matrix **X** and matrix **Y**. Regularization factor *λ* = 0.5, and entropic regularization term is specified, to control the generation of one-to-many pairings:

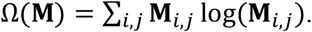

Sinkhorn-Knopp matrix scaling algorithm[29] is adopted to solve the above optimal transport problem.

After working out the optimal matrix **M** which stores matching weight between each control and case cell, for each case cell, its matched control cell with the highest weight is selected as its final match, denoted as 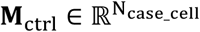:

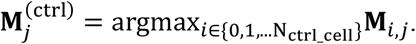

Then the matched control expression matrix is acquired by re-ordering cells:

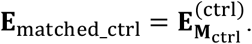

### DCAF-criterion

#### Removal of genes that remain silent

According to DCAF-criterion, before assessing the perturbation prediction accuracy of a method, genes that remain silent throughout perturbation for each cell are filtered out, to remove their disturbance to the assessment. To establish the criteria of filtration, a threshold for preprocessed expression of each test dataset is specified through manually observing its expression distribution. The established threshold shall separate zero inflation part and negative binomial part of the complete distribution. The threshold value thrsh is then applied onto matched control and case expression to get a mask 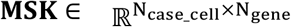:

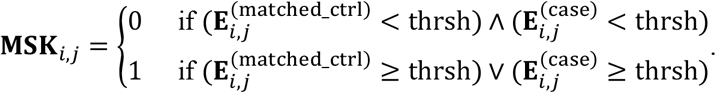

To avoid considering genes that remain silent when measuring model accuracy, **MSK** is equally applied onto (matched) control, case and predicted expression matrix. For these cells, three vectors are respectively generated to save control, case, and predicted case expression of genes that are activated, silenced, or remain expressed. Remained genes can be different across single cells.

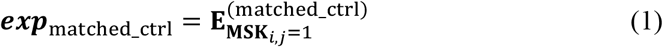

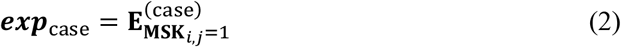

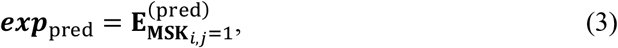

where 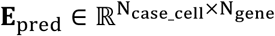 is the predicted case transcriptomics after normalization and log1p transformation.

#### R^**2**^ of log fold change

To measure model accuracy on genes do not remain silenced, DCAF-criterion measures a method’s performance, by comparing the ground truth and its predicted transcriptomics after perturbation on each single cell and averages the results across all cells. To calculate this index, DCAF-criterion first measures the log fold change of predicted and true transcriptomics then solves the accuracy by comparing them. For test cells, according to equations (1), (2) and (3), ***exp***_matched_ctrl_, ***exp***_case_ and ***exp***_pred_ store control, case and predicted transcriptomics in vectors. Built on the three vectors, we first calculate the true log fold change value ***LFC***_true_, and the predicted log fold change value ***LFC***_pred_:

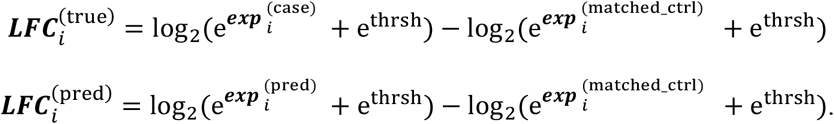

Then

R^2^ of log fold change (for genes do not remain silenced) = R^2^(***LFC***_pred_, ***LFC***_true_), where R^2^(***X, Y***) is the coefficient of determination[51] of vectors ***X*** and ***Y***.

#### MSE of log fold change

To calculate this matrix for genes do not remain silenced, we first work out the log fold change of predicted and true gene expression, ***LFC***_pred_ and ***LFC***_true_, as described in R^2^ of log fold change.

Then:

MSE of log fold change (for genes do not remain silenced) = MSE(***LFC***_pred_, ***LFC***_true_), where MSE(***X, Y***) is the mean squared error[52] of vectors ***X*** and ***Y***.

#### MSE and MAE

For genes that remain silenced, MSE and MAE are used by DCAF-criterion to measure model accuracy.

To start with, three vectors are respectively generated to save control, case, and predicted case expression of genes that remain silenced. Remaining silenced genes can be different across single cells.

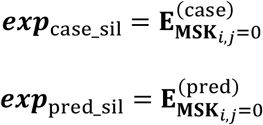

Then:

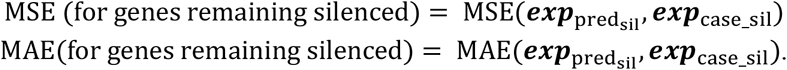

### DCAF-model

#### Model Structure

Different from custom generative structure that projects predicted case gene embeddings to case gene expressions at its top layers, JEPA structure does not involve this reconstruction process. Instead, in the first phase, it represents case transcriptomics as case gene embeddings by encoder, and the same encoder and a predictor are trained together, to minimize the distance between predicted and true case gene embeddings. Later in the second phase, the necessary reconstruction process is implemented by training the reconstructor structure alone, to project predicted case gene embeddings to case transcriptomics.

Specifically, by JEPA structure, control gene expression 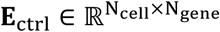 are encoded to 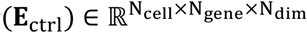, and the case cells are encoded to 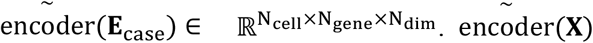 is the estimated moving average[53] of encoder(**X**). We use 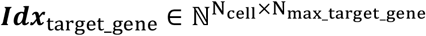 to denote targeted genes of each cell, where N_max_target_gene_ represents max. no. of perturbed genes of each cell in the dataset. For each perturbed cell, the column indices of its perturbed genes on **E**_ctrl_ and **E**_case_ are inserted as elements on respective rows ***Idx***_target_gene_. On each row, -9999 is inserted to each column that are not inserted with indices. encoder(**E**_ctrl_) and ***Idx***_target_gene_ are then injected into a perturbation encoder pert(**X, *Y***), to encode the perturbation information, and this perturbation information and encoder(**E**_ctrl_) are together fed into a 3-layer transformer predictor(**X, Y**), to generate predicted case gene embeddings predictor(encoder(**E**_ctrl_), 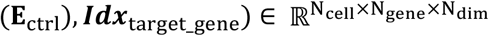. The embeddings are measured in comparison to true case gene embeddings 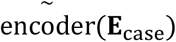, for the loss minimization:

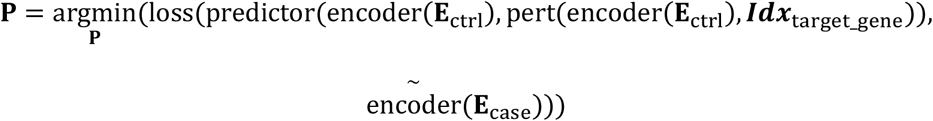

Thereinto, **P** denotes training parameters of the JEPA structure, encoder(**X**) is adapted from scGPT[18], with LoRA structure[33] added to Q, K, V and O projections for finetuning, using rank of 4. During training of JEPA, pretrained parameters of encoder remain frozen and all other parameters including LoRA and predictor parameters are open for training.

By perturbation encoder pert(**X, Y**), for each cell, the control embedding of perturbed gene is adopted as the universal embedding across all genes. For any cell, if over 1 gene is perturbed, the perturbed gene embeddings are average pooled to generate the embedding for propagation:

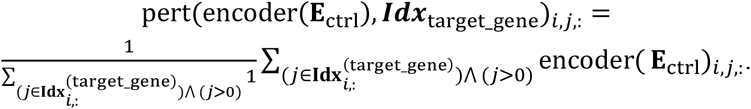

By predictor predictor(**X, Y**) , both **X** and **Y** inputs are first projected from encoder dimension to 64, through linear probes respectively applied onto the last dimension of two matrices. The main skeleton of the predictor is a 3-layer Transformer architecture[54]. In the first layer, the projected **X** is adopted as the Q and K and the projected **Y** is adopted as V term for multi-head cross-attention with head no. of 8. After the attention operation, the 64-dimension **X** is added back to attention result, in Add & Normalization process. Then output of the first layer is fed to the next two transformer layers for multi-head self-attention, with structure parameters remaining the same as the first layer. After the 3^rd^ layer, the last dimension is projected back to encoder dimension. During the training of JEPA, all parameters of the predictor are open for training.

By reconstructor structure, both cross-gene and cross-representation dimension operation are applied on predicted case gene representation, to project the representation to gene expression. Cross-gene operation implements the following process:

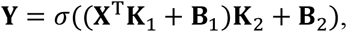

where 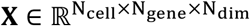 denotes the output of predictor, **X**^T^ swaps the last two dimension, 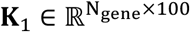, 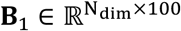, 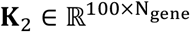, 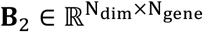 and *σ*(**X**) denotes the activation function Leaky ReLU, with dropout applied with dropout rate of 0.1. Both **K**_1_ and **B**_1_ are frozen while **K**_2_ and **B**_2_ are open for training. By cross-representation dimension operation:

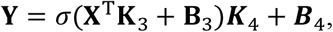

where 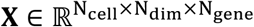 denotes the output of cross-gene operation, **X**^T^ swaps the last two dimension, 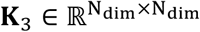, 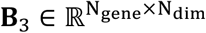, 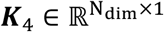, 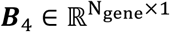 and*σ*(**X**) denotes the activation function Leaky ReLU, with dropout applied with dropout rate of 0.1. When training the reconstructor, previously finetuned encoder and trained predictor are both frozen, while all reconstructor parameters are open for training.

#### Data Preprocessing

Before using perturbation data to train our model, Quality Control (QC) is first implemented considering UMI counts of each cell. Python function scanpy.pp.filter_cells were used to implement this process, with minimum counts of UMI assigned as 200. Next, genes not included in scGPT’s gene dictionary are removed. The expression matrix is normalized based on each cell’s total counts, using scanpy.pp.normalize_total, before being transformed by scanpy.pp.log1p. Then if over 5000 genes are included in expression matrix, only top 5000 Highly Variable Genes (HVG) are kept, using scanpy.pp.highly_variable_genes. Additionally, genes that are not highly variable yet being perturbed are added back to the expression matrix. As required by our model, a case cell is removed if any one of its target genes falls out of gene list of expression matrix. Finally on this processed matrix, if zero value proportion reaches over 0.8, genes with top 800 expressed cells are re-ordered, to be uniformed distributed along gene axis of expression matrix. This is to induce a more uniform distribution of transcriptomics fed into LLM through data loader. Because when training, validating or testing our proposed model, instead of being truncated, transcriptomics is randomly sampled in some scenarios and continuously sliced in others, a more uniform distribution brings a more uniform performance of LLM when representing each cell.

For a more balanced training corpus across different targeted genes, necessary down-sampling is implemented on preprocessed data, before feeding them to our model. When training our model based on cell-line dataset, 50 cells are randomly sampled for each group of case cells with the same target gene or the same set of target genes. Upon a group with less than 50 cells, all cells remain. When training our model based on multi cell type data, 100 cells are randomly sampled for each group of case cells with the same annotation type and same target gene or genes. Upon a group with less than 100 cells, all cells are kept. Tested cells are not down sampled.

### Data loader for Training and Validation

The training corpus is then divided to be respectively fed to training and validating data loaders. If the model is trained to simulate perturbations on novel genes, 82% of target genes are randomly selected, and their corresponding case cells are fed to training data loader. And the left cells are fed to validation data loader. If the model is trained to simulate perturbations on novel cell types, 82% of cells are randomly selected as training cells, and the left cells are used for validation.

Either when training JEPA or when training reconstructor, in training data loaders, all preprocessed cells are shuffled and sent to our model in different batches. While for validating data loaders, if over 2000 cells are to be fed, only 2000 of them are randomly sampled to save time consumption of validation. However, after the down-sampling process, datasets are then re-adjusted in different ways comparing data loader of JEPA and data loader of reconstructor. When training the JEPA structure, a different random 1200 genes are sampled for each training and validating batch, to maintain the cost-effectiveness during attention calculation in the LLM encoder. When training the reconstructor, all genes after preprocessing remain in either training or validating data loaders.

#### Training Methods

Different parts of the model are frozen during JEPA training and reconstructor training sessions. When JEPA is trained, the pretrained parameters of LLM are frozen, with LoRA parameters of LLM and all parameters of predictor open for updates. While reconstructor is trained, both pretrained and LoRA parameters of LLM, as well as predictor parameters are frozen, leaving only reconstructor parameters open for updates.

Also, during the training of JEPA, an exponential moving average of control expression’s encoder is applied onto case transcriptomics, with decay value of 0.995. Specific implementation is according to Python class: ExponentialMovingAverage provided by package torch_ema.

Either when training JEPA or when training reconstructor, Adam optimizer is used, with learning rate set as 1 e-3, term added to the denominator set as 1 e-4, and weight decay set as 5 e-4. Step learning rate is adopted at the end of each epoch, with multiplicative factor equal to 0.5. Automatic mixed precision functionality of PyTorch is enabled, with gradient scaling adopted. Except from these mentioned hyperparameters, all unspecified hyperparameters adopt default values provided by PyTorch, in version of 2.0.0+cu118, through classes torch.optim.Adam, torch.optim.lr_scheduler.StepLR, torch.autocast and torch.cuda.amp.GradScaler. Model’s performance on validation is measured at the end of each epoch, and early stop is triggered when the model stops improving for 5 consecutive epochs, or when the model is trained for at least 20 epochs.

### Experiment and Data Analysis Methods

#### Baselines for Benchmarking DCAF-model in Pre- and Post Perturbation Status Matching

For baseline of random matching, all pre- and post-perturbation cells are randomly paired to each other. For baseline of random matching within cell types, each case cell is randomly matched with a control cell that has the same cell type. CINEMA-OT was implemented according to its published article[22].

### Data Preprocessing for Benchmarking DCAF-model in Pre- and Post Perturbation Status Matching

For all matching methods compared, datasets were first processed following the same procedure as described in Data Preprocessing of DCAF-model. The training and testing case cells were then concatenated to generate case cells for matching. All the preprocessed control cells were then used to match case cells.

### Synthetic Data Generation for Comparing Accuracy Matrices to Measure Response

To study DCAF-criterion’s response to different levels of model accuracy, we first generate simulation datasets with different known level of errors. Specifically, control cells were first produced to obey zero-inflated binomial distribution, with zero expression proportion of 83%. The shape parameters of negative binomial distribution include number of successes r = 2 and probability of single success p = 0.1. The distribution was moved 10 units in the right direction of its original position to be closer to actual distributions. Specifically, to implement the above process, we denote 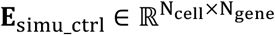 to represent UMI count of control cells, where each element X of **E**_simu_ctrl_ follows:

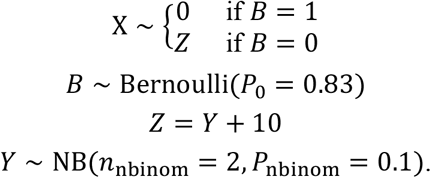

Then control cell expression was sum normalized with sum of each cell’s expression equal to 1000.

When generating case cells, elements of expression matrix were swapped, shuffled, or remain unchanged, to keep the ratio of activated, silenced, remaining expressed and remaining silenced element consistent with published literature[38][39][35][27][40][36]. Specifically, 47% of positive expressions were swapped with zero expressions. The positive expressions left un-swapped were automatically shuffled to mimic change of genes remaining expressed. By this way, 8% of genes are activated, 8% of genes are silenced, 9% of genes remain expressed, while the left 75% remains silenced throughout the perturbation.

To simulate different model performance, an exponentially changing standard deviation of Gaussian noise was applied onto overall case data **E**_simu_case_, with specific standard deviation *std* ranging from 0.02, 0.05, 0.1, 0.2, 0.4 to 0.8:

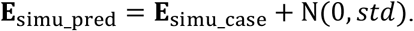

#### Synthetic Data Generation for Comparing Robustness of Accuracy Matrices

To study robustness of DCAF-criterion against different model outputs, we generate simulation dataset with different known levels of disturbances. First, control and case gene expressions were generated following the same procedure as described in Synthetic Data Generation for Comparing Matrices to Measure Response. Then Gaussian noise with various standard deviation were injected into zero expressions part only, to simulate disturbances that cause different model output yet not different model error levels. The values of standard deviation *std* include 0.02, 0.05, 0.1, 0.2, 0.4 and 0.8 (Fig. 3e, supp). To formulate this process:

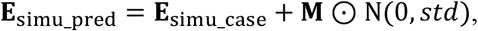

Where 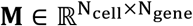, and

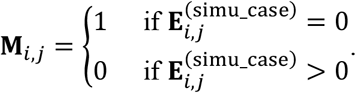

#### Baselines for Benchmarking DCAF-model

Benchmarking methods of GEARS[17] and scGPT[18] were implemented according to their original publication. Default parameters of their code in the public repositories were used for training and testing.

Linear method was adopted by fitting the elements of a gene-wise multiplication matrix 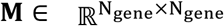, to minimize the training loss:

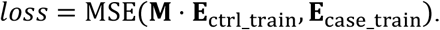

Hyperparameters specified in Training Methods of DCAF-model were used for training process. Then the solved matrix **M** is used to predict case gene expression of test cells:

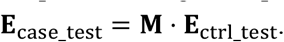

Details of mean baseline vary slightly on dataset with single and multiple cell types. On cell-line data with one single cell type, to predict post-perturbation status on novel target genes, an average gene expression across all training cells is used, as the predicted expression of each tested cell. While in scenarios of predicting results on novel cell types, as cells of the same cell type are targeted with different genes, only training cells with same perturbed gene(s) are calculated average gene expression, to predict expression of different cell types with the same target gene.

#### Data Preprocessing for Benchmarking DCAF-model

On mean baseline, linear baseline, and DCAF-model, same data preprocessing was applied on datasets before testing their accuracy, according to the description in Data Preprocessing of DCAF-model. When studying accuracy on novel gene perturbations of these methods, 85% of preprocessed cells were used for training, and 15% were used for testing. Specifically, training and testing cells are targeted with totally different genes or groups of genes during multi-gene perturbation. When studying models’ accuracy on novel cell-type perturbations, a random 70% of cell types were used for training, and the left cell types were used for testing. The training data acquired by above methods were then split into training and validation dataset before feeding to respective data loaders, described in DCAF-guided Model session. In testing data loaders, cells were processed in the same way as in training and validation data loaders of DCAF-model’s reconstructor, as described in DCAF-model.

Data Preprocessing for GEARS was implemented as specified in its article[17]. For target genes which correspond to only one case cell, the corresponding case cells are filtered out, to avoid the program from breaking off.

Data Preprocessing for scGPT was implemented according to its article[18]. We further removed case cells with target gene(s) falling out of gene list of expression matrix. This operation is implemented to avoid the program from breaking off.

In Frangieh dataset, only cancer cells co-cultured with T cells were used in either training or testing, for all methods including GEARS, scGPT, linear baseline, mean baseline and DCAF-model. This was implemented to better understand the performance of each model, upon a consistent micro-environment applied on all control and case cells.

#### Data Preprocessing for Studying DCAF-model Using Human Brain Organoid Data

When studying DCAF-model’s capability in transferring capability of perturbation simulation from seen to unseen genes in Human brain organoid data, 70% of perturbed targets and its corresponding case cells were used as training dataset, 15% were used as validation dataset and 15% were used as testing dataset. When studying the model’s capability in transferring to unseen cell types, across different transfer studies, each one of cell types including Astrocyte, Excitatory, Inhibitory, NPCs and OPC was separated from other cell types, and its corresponding cells were used as testing data. A random 82% of left cells were used as training datasets, and the left 18% of cells were used for validation.

In testing data loaders, cells were processed in the same way as in training and validation data loaders for DCAF-guided reconstructor, as described in DCAF-model.

#### Data Preprocessing for Studying DCAF-model Using Adult Mouse Brain Data

When studying DCAF-model’s capability in transferring perturbation simulation capability to unseen cell types using adult Mouse brain dataset, a random 70% of cell types were used for training, and the left cell types were used for testing. The training data was then split into training and validation dataset before feeding to respective data loaders, described in DCAF-model session. After the model training, the trained model was applied onto MOp region of Allen’s Mouse brain. All cell types were used without filtration as test corpus.

In testing data loaders, cells were processed in the same way as in training and validation data loaders for DCAF-model reconstructor, as described in DCAF-model.

#### Analysis Metrices

Following metrices are used in our research to quantify dataset’s characteristics, measurements and methods’ accuracy.

#### Mean KL divergence

To define this matrix, we introduce 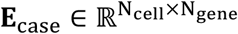 to denote preprocessed gene expression of case cells, where each row represents a single cell and each column represents a gene. Then we use 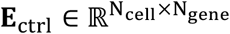 to denote preprocessed gene expression of control cells, with each row describing a single cell expression matched to a cell in ***E***_case_ with the same row index, and each column representing a gene, ordered in the same position as in **E**_ctrl_. Then mean KL divergence matrix calculates the average divergence of matched cells, by:

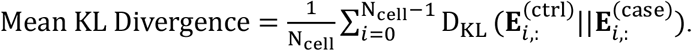

Thereinto, D_KL_(***p***||***q***) denotes the Kullback-Leibler divergence[50] measurement between vector ***p*** and vector ***q***.

#### Single-cell Level Systema Accuracy

To adapt the centroid accuracy introduced by Systema framework[24] on our single cell level accuracy assessment, we use the centroid of perturbed cells as reference and draw vectors joining this reference and each of predicted and true case cells. To be consistent with the original research of Systema, Pearson correlation was adopted to measure the difference between predicted and true transcriptomics. To specifically define this measurement, we further introduce 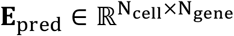 to represent predicted normalized and transformed gene expression of case cells. Then:

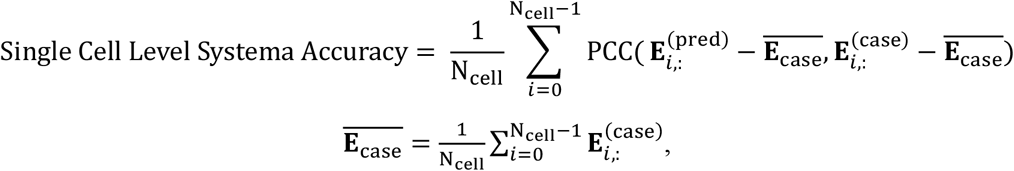

where PCC(***p, q***) denotes Pearson Correlation Coefficient[55] of vector ***p*** and ***q***.

#### Up-regulation F1 Score

Up-regulation F1 Score quantifies a given method’s capability in predicting the up-regulated genes on a specific dataset, by calculating the harmonic mean of its precision and recall score on given testing scenarios. Specifically, we first calculate the true log fold change value 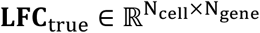, and the predicted log fold change value 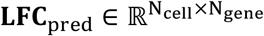, where:

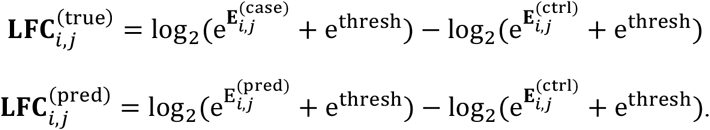

We then introduce 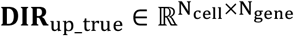 to represent the discriminative result by telling if each gene in each cell has actually gone through a log fold change exceeding threshold of 2, and 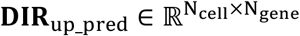 to represent the similar meaning, except by being predicted. Specifically:

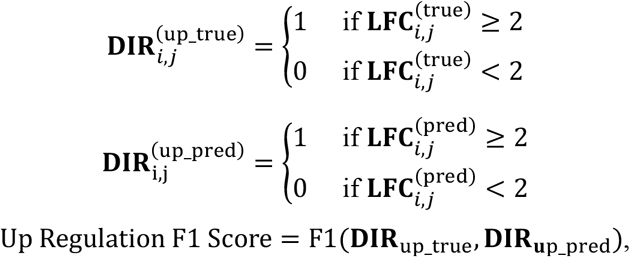

where**F1**(**True, Pred**) is the F1-score of true and predicted array of binary classification results.

When remain-silenced genes are not considered when calculating this matrix, positions of elements representing those genes in specific cells were calculated according to DCAF-criterion. And those elements were removed during the calculation of F1-score as described in the same section.

#### Unsignificant-regulation F1 Score

Unsignificant-regulation F1 Score quantifies a given method’s capability in predicting the unsignificant-regulated genes on a specific dataset, by harmonic mean of its precision and recall score on the given testing scenario. Specifically, after calculating the true log fold change value ***LFC***_true_, and the predicted log fold change value ***LFC***_pred_ described in Up-regulation F1 Score, we introduce 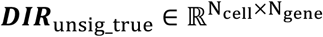 to represent the discriminative result by telling if each gene in each cell was actually insignificantly-regulated with log fold change falling between -2 and 2, and 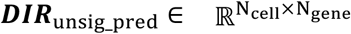 to represent the similar meaning, except by prediction result. Specifically:

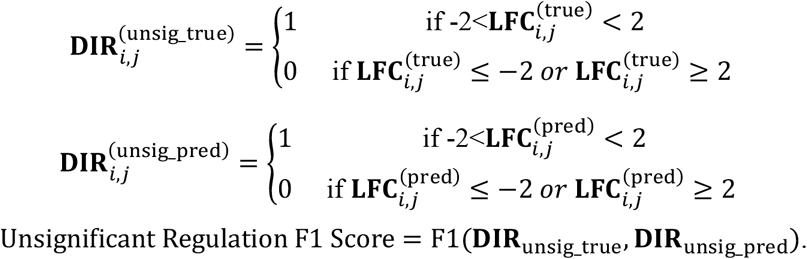

When remain-silenced genes are not considered when calculating this matrix, positions of elements representing those genes in specific cells were calculated according to DCAF-criterion of accuracy measurement. And those elements were removed during the calculation of F1-score as described in the same section.

#### Down-regulation F1 score

Down-regulation F1 Score quantifies a given method’s capability in predicting the down-regulated genes on a specific dataset, by harmonic mean of its precision and recall on the given testing scenario. Specifically, after calculating the true log fold change value **LFC**_true_, and the predicted log fold change value **LFC**_pred_ described in Up-regulation F1 Score, we introduce 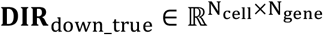 to represent the discriminative result by telling if each gene in each cell was actually down-regulated, with log fold change falling under -2. And we introduce 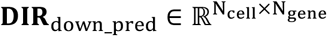 to represent the similar meaning, except by prediction result. Specifically:

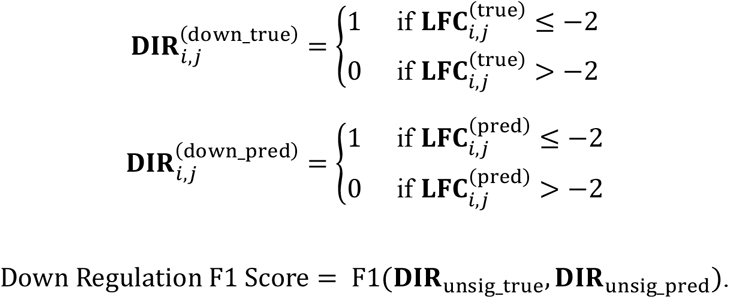

When remain-silenced genes are not considered when calculating this matrix, positions of elements representing those genes in specific cells were calculated according to DCAF-criterion. And those elements were removed during the calculation of F1-score as described in the same section.

#### Cell type-wise cosine similarity

Only control cells of Human brain organoid dataset were used during this measurement. The cells were preprocessed by filtration with UMI counts less than 200, using scanpy.pp.filter_cells, and by normalization with scanpy.pp.normalize_total, and eventually by transformation with scanpy.pp.log1p. Then only top 5000 HVG remained by HVG selection using scanpy.pp.highly_variable_genes. To accelerate consequent computation, only 100 cells were randomly selected for each cell type with over 100 cells, and all cells remained selected when it came to cell types with less than this amount. Then based on the down sampled cells, an average cell for each cell type was calculated by average pooling within the group. To formulate this process, we introduce 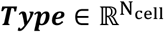 to denote cell types of each preprocessed and down sampled cell, 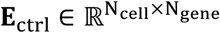 to store their gene expressions, as well as all-ones vector 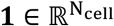. We have the following for a given cell type *x*:

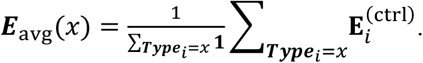

Then for given cell type *x* and *y*, their:

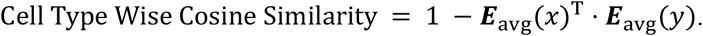

#### Systematic similarity

We transfer the concept of systematic variation across different perturbed genes from Systema[24] into our study to understand the difficulty to predict perturbation effect on novel cell types. Two core differences are present between the study of Systema and ours, and they bring alterations from original systematic variation to our systematic similarity. First, our model is required to transfer from novel target genes generalization to novel cell type generalization. Second, instead of measuring a single variation of the complete dataset, we try to understand how different train-test split methods lead to different variations.

To inherit the thought of systematic variation and to deal with the differences, we quantified the similarity between center of trained cell types and the test cell type for each way of split. Specifically, case cells of Human brain organoid dataset were used and preprocessed under the same procedure as described in Cell type-wise cosine similarity. Suppose cell type *c* is left out as the test cell type and all other cell types are used for training, we first calculate average cell ***E***_avg_(*c*) for *c* also described in the above section. To denote the average status of the training cell types, we average across cell types other than *c*:

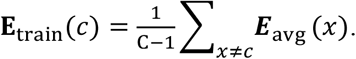

Thereinto C denotes total number of cell types. Then for given cell type *c*,

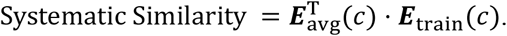

#### Normalized Accuracy

Normalized accuracy is calculated as the normalization result between accuracy of tested method and a baseline method[56][57]. We denote 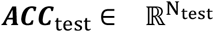 and 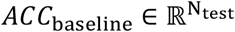 to respectively represent tested and baseline method accuracy on N_test_ number of scenarios. Then the normalized accuracy is calculated as:

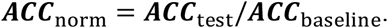

In this study, we measured the normalization of R^2^ of log change accuracy between DCAF-model as tested method, and mean baseline as reference method, on each cell type.

#### Systematic Dispersion

We propose the concept of Systematic Dispersion to depict level of heterogeneity for each cell type, by calculating averaged pairwise distances across cells. Specifically, tested case cells of each cell type of Human brain organoid dataset were adopted, and they were preprocessed according to Data Preprocessing of DCAF-model. Top 50 components are selected in PCA of expression data, by using scanpy.tl.pca. And Euclidean distance was calculated between PCA representations of every two cells within a certain cell type, before being averaged to systematic dispersion result. To formulate this process, we denote 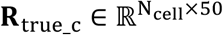 to represent PCA representation of ground truth post-perturbation expression of tested cell type *c*. Then its:

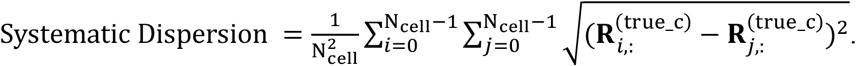

## Data availability

Gene Expression Omnibus accession numbers used for adult Mouse brain[35] and *in utero* Mouse brain[34] are GSE236519, and GSE157977. Human brain organoid[36] dataset is available on Zenodo at https://zenodo.org/records/7083558. Data from Norman et al.[39], Frangieh et al.[38], and Replogle et al.[27] are available at https://projects.sanderlab.org/scperturb/datavzrd/scPerturb_vzrd_v1/dataset_info/index_1.html. H1 hESCS[40] dataset was downloaded from https://github.com/ArcInstitute/arc-virtual-cell-atlas/blob/main/virtual-cell-challenge/README.md. The data from Tian et al.[37] are available at http://www.perturbase.cn/repository.

## Code availability

Scripts for this study are publicly available at: https://github.com/BGIResearch/DCAF.

## Acknowledgements

We acknowledge DCS Genpilot platform (genpilot.dcs.cloud) for providing convenient ways for research and analysis. We appreciate Dr. Jianhua Yin and Dr. Weikai Wang for their insightful discussions on perturbations. We would like to express our sincere gratitude to Dr. Sheng Zhang, Dr. Rongjiang Zhao, Dr. Dong Zhang, Ms. Zhe Li, Dr. Feng Ren, and Dr. Yipeng Wang, for their professional support and thoughtful actionable guidance throughout this research. We also thank Mr. Zirong Zhao, Mr. Yangbang Shang and Ms. Wurenqimuge for their explorative efforts in method development during the initial stage of this project.

## Author contributions

T.X. and S. F. conceived this study. T.X designed DCAF-data and DCAF-model. She also worked with C.G. in designing DCAF-criterion. T.X. performed most of method development, research and analysis, and L.C. performed a substantial proportion of research and analysis in DCAF-model explainability. T.X and N.C wrote the manuscript. S.F supervised this research. All authors provide constructive discussions that lead to design of the method and research.

## References

[1] Kitano, H.: Systems biology: a brief overview. science 295(5560), 1662–1664 (2002).

[2] Shi, H., Doench, J.G., Chi, H.: Crispr screens for functional interrogation of immunity. Nature Reviews Immunology 23(6), 363–380 (2023).

[3] Jinek, M., Chylinski, K., Fonfara, I., Hauer, M., Doudna, J.A., Charpentier, E.: A programmable dual-rna–guided dna endonuclease in adaptive bacterial immunity. science 337(6096), 816–821 (2012).

[4] Dixit, A., Parnas, O., Li, B., Chen, J., Fulco, C.P., Jerby-Arnon, L., Marjanovic, N.D., Dionne, D., Burks, T., Raychowdhury, R., et al.: Perturb-seq: dissecting molecular circuits with scalable single-cell rna profiling of pooled genetic screens. cell 167(7), 1853–1866 (2016).

[5] Zhang, Y., Huang, J., Li, Y., Liu, T., Ren, L., Huang, J., Lin, H., Feng, J., Zhang, Y.: Charting the computational landscape of single-cell genetic perturbation. Journal of Advanced Research (2026).

[6] Gavriilidis, G.I., Vasileiou, V., Orfanou, A., Ishaque, N., Psomopoulos, F.: A mini-review on perturbation modelling across single-cell omic modalities. Computational and Structural Biotechnology Journal (2024).

[7] Menon, A.V., Song, B., Chao, L., Sriram, D., Chansky, P., Bakshi, I., Ulianova, J., Li, W.: Unraveling the future of genomics: Crispr, single-cell omics, and the applications in cancer and immunology. Frontiers in genome editing 7, 1565387(2025).

[8] Vanhaelen, Q., Aliper, A., Zhavoronkov, A.: A comparative review of compu-tational methods for pathway perturbation analysis: dynamical and topological perspectives. Molecular BioSystems 13(9), 1692–1704 (2017).

[9] Aon, M.A., Cortassa, S.: Systems biology of the fluxome. Processes 3(3), 607–618 (2015).

[10] Tomar, N., De, R.K.: Comparing methods for metabolic network analysis and an application to metabolic engineering. Gene 521(1), 1–14 (2013).

[11] Rukhlenko, O.S., Halasz, M., Rauch, N., Zhernovkov, V., Prince, T., Wynne, K., Maher, S., Kashdan, E., MacLeod, K., Carragher, N.O., et al.: Control of cell state transitions. Nature 609(7929), 975–985 (2022).

[12] Barabasi, A.-L., Oltvai, Z.N.: Network biology: understanding the cell’s functional organization. Nature reviews genetics 5(2), 101–113 (2004).

[13] Bruggeman, F.J., Westerhoff, H.V.: The nature of systems biology. TRENDS in Microbiology 15(1), 45–50 (2007).

[14] Tomar, N., K De, R.: A comprehensive view on metabolic pathway analysis methodologies. Current Bioinformatics 9(3), 295–305 (2014).

[15] Papoutsakis, E.T.: Equations and calculations for fermentations of butyric acid bacteria. Biotechnology and bioengineering 26(2), 174–187 (1984).

[16] Watson, M.: Metabolic maps for the Apple II. Portland Press Ltd. (1984).

[17] Roohani, Y., Huang, K., Leskovec, J.: Predicting transcriptional outcomes of novel multigene perturbations with gears. Nature Biotechnology 42(6), 927–935 (2024).

[18] Cui, H., Wang, C., Maan, H., Pang, K., Luo, F., Duan, N., Wang, B.: scgpt: toward building a foundation model for single-cell multi-omics using generative ai. Nature methods 21(8), 1470– 1480 (2024).

[19] Adduri, A.K., Gautam, D., Bevilacqua, B., Imran, A., Shah, R., Naghipourfar, M., Teyssier, N., Ilango, R., Nagaraj, S., Dong, M., et al.: Predicting cellular responses to perturbation across diverse contexts with state. BioRxiv, 2025–06 (2025).

[20] Behjati Ardakani, F., Kattler, K., Heinen, T., Schmidt, F., Feuerborn, D., Gasparoni, G., Lepikhov, K., Nell, P., Hengstler, J., Walter, J., Schulz, M.H.: Prediction of single-cell gene expression for transcription factor analysis. GigaScience 9(11) (2020).

[21] Bunne, C., Stark, S.G., Gut, G., Del Castillo, J.S., Levesque, M., Lehmann, K.-V., Pelkmans, L., Krause, A., Rätsch, G.: Learning single-cell perturbation responses using neural optimal transport. Nature methods 20(11), 1759–1768 (2023).

[22] Dong, M., Wang, B., Wei, J. O., Fonseca, A.H., Perry, C.J., Frey, A., Ouerghi, F., Foxman, E.F., Ishizuka, J.J., Dhodapkar, R.M., et al.: Causal identification of single-cell experimental perturbation effects with cinema-ot. Nature methods 20(11), 1769–1779 (2023).

[23] Li, C., Gao, H., She, Y., Bian, H., Chen, Q., Liu, K., Wei, L., Zhang, X.: Benchmarking ai models for in silico gene perturbation of cells. bioRxiv. 2024a (2024).

[24] Viñas Toráe, R., Wiatrak, M., Piran, Z., Fan, S., Jiang, L., Teichmann, S.A., Nitzan, M., Brbić, M.: Systema: a framework for evaluating genetic perturbation response prediction beyond systematic variation. Nature Biotechnology, 1–10 (2025).

[25] Wong, D.R., Hill, A.S., Moccia, R.: Simple controls exceed best deep learning algorithms and reveal foundation model effectiveness for predicting genetic perturbations. Bioinformatics 41(6), 317 (2025).

[26] Gretton, A., Borgwardt, K.M., Rasch, M.J., Schölkopf, B., Smola, A.: A kernel two-sample test. The journal of machine learning research 13, 723–773 (2012).

[27] Replogle, J.M., Saunders, R.A., Pogson, A.N., Hussmann, J.A., Lenail, A., Guna, A., Mascibroda, L., Wagner, E.J., Adelman, K., Lithwick-Yanai, G., et al.: Mapping informationrich genotype-phenotype landscapes with genome-scale perturb-seq. Cell 185(14), 2559–2575 (2022).

[28] Lotfollahi, M., Wolf, F.A., Theis, F.J.: scgen predicts single-cell perturbation responses. Nature methods 16(8), 715–721 (2019).

[29] Cuturi, M.: Sinkhorn distances: Lightspeed computation of optimal transport. Advances in neural information processing systems 26 (2013).

[30] Wang, Z., Zhan, Q., Yang, S., Zhou, Z., Kan, M., Zhai, T., Shen, L.: An interpretable graphregularized optimal transport framework for diagonal single-cell integrative analysis. GigaScience 15, 012 (2026).

[31] Assran, M., Duval, Q., Misra, I., Bojanowski, P., Vincent, P., Rabbat, M., LeCun, Y., Ballas, N.: Self-supervised learning from images with a joint-embedding predictive architecture. In: 2023 IEEE/CVF Conference on Computer Vision and Pattern Recognition (CVPR), pp. 15619–15629 (2023). IEEE.

[32] Bardes, A., Garrido, Q., Ponce, J., Chen, X., Rabbat, M., LeCun, Y., Assran, M., Ballas, N.: Revisiting feature prediction for learning visual representations from video. arXiv preprint arXiv:2404.08471 (2024).

[33] Hu, E.J., Shen, Y., Wallis, P., Allen-Zhu, Z., Li, Y., Wang, S., Wang, L., Chen, W., et al.: Lora: Low-rank adaptation of large language models. Iclr 1(2), 3 (2022).

[34] Zheng, X., Wu, B., Liu, Y., Simmons, S.K., Kim, K., Clarke, G.S., Ashiq, A., Park, J., Li, J., Wang, Z., et al.: Massively parallel in vivo perturb-seq reveals cell-type-specific transcriptional networks in cortical development. Cell 187(13), 3236–3248 (2024).

[35] Santinha, A.J., Klingler, E., Kuhn, M., Farouni, R., Lagler, S., Kalamakis, G., Lischetti, U., Jabaudon, D., Platt, R.J.: Transcriptional linkage analysis with in vivo aav-perturb-seq. Nature 622(7982), 367–375 (2023).

[36] Li, C., Fleck, J.S., Martins-Costa, C., Burkard, T.R., Themann, J., Stuempflen, M., Peer, A.M., Vertesy, À., Littleboy, J.B., Esk, C., et al.: Single-cell brain organoid screening identifies developmental defects in autism. Nature 621(7978), 46 (2023).

[37] Tian, R., Abarientos, A., Hong, J., Hashemi, S.H., Yan, R., Dräger, N., Leng, K., Nalls, M.A., Singleton, A.B., Xu, K., et al.: Genome-wide crispri/a screens in human neurons link lysosomal failure to ferroptosis. Nature neuroscience 24(7), 1020–1034 (2021).

[38] Frangieh, C.J., Melms, J.C., Thakore, P.I., Geiger-Schuller, K.R., Ho, P., Luoma, A.M., Cleary, B., Jerby-Arnon, L., Malu, S., Cuoco, M.S., et al.: Multimodal pooled perturb-cite-seq screens in patient models define mechanisms of cancer immune evasion. Nature genetics 53(3), 332– 341 (2021).

[39] Norman, T.M., Horlbeck, M.A., Replogle, J.M., Ge, A.Y., Xu, A., Jost, M., Gilbert, L.A., Weissman, J.S.: Exploring genetic interaction manifolds constructed from rich single-cell phenotypes. Science 365(6455), 786–793 (2019).

[40] Roohani, Y.H., Dobin, A., Teyssier, N., Eraslan, B., Hua, T.J., Tung, P.-Y., Bounds, L.R., Adduri, A., Woodrow, A., Plosky, B.S., et al.: Behind the data of the virtual cell challenge. Arc Institute, August 2025a. URL https://arcinstitute.org/news/behind-the-data-virtual-cellchallenge (2025).

[41] Wang, Q., Ding, S.-L., Li, Y., Royall, J., Feng, D., Lesnar, P., Graddis, N., Naeemi, M., Facer, B., Ho, A., et al.: The allen mouse brain common coordinate framework: a 3d reference atlas. Cell 181(4), 936–953 (2020).

[42] Stark, K.L., Xu, B., Bagchi, A., Lai, W.-S., Liu, H., Hsu, R., Wan, X., Pavlidis, P., ills, A.A., Karayiorgou, M., et al.: Altered brain microrna biogenesis contributes o phenotypic deficits in a 22q11-deletion mouse model. Nature genetics 40(6), 751–760 (2008).

[43] Schofield, C.M., Hsu, R., Barker, A.J., Gertz, C.C., Blelloch, R., Ullian, E.M.: Monoallelic deletion of the microrna biogenesis gene dgcr8 produces deficits in the development of excitatory synaptic transmission in the prefrontal cortex. Neural development 6(1), 11 (2011).

[44] JnBaptiste, C.K., Gurtan, A.M., Thai, K.K., Lu, V., Bhutkar, A., Su, M.-J., Rotem, A., Jacks, T., Sharp, P.A.: Dicer loss and recovery induce an oncogenic switch driven by transcriptional activation of the oncofetal imp1–3 family. Genes & development 31(7), 674–687 (2017).

[45] Hsu, R., Schofield, C.M., Cruz, C.G.D., Jones-Davis, D.M., Blelloch, R., Ullian, E.M.: Loss of micrornas in pyramidal neurons leads to specific changes in inhibitory synaptic transmission in the prefrontal cortex. Molecular and Cellular Neuroscience 50(3-4), 283–292 (2012).

[46] Daswani, R., Gilardi, C., Soutschek, M., Nanda, P., Weiss, K., Bicker, S., Fiore, R., Dieterich, C., Germain, P.-L., Winterer, J., et al.: Microrna-138 controls hippocampal interneuron function and short-term memory in mice. Elife 11, 74056 (2022).

[47] He, M., Liu, Y., Wang, X., Zhang, M.Q., Hannon, G.J., Huang, Z.J.: Cell-type-based analysis of microrna profiles in the mouse brain. Neuron 73(1), 35–48 (2012).

[48] Delvendahl, I., Daswani, R., Winterer, J., Germain, P.-L., Uhr, N.M., Schratt, G., Müller, M.: Microrna-138-5p suppresses excitatory synaptic strength at the cerebellar input layer. The Journal of Physiology 603(10), 3161–3179 (2025).

[49] Oliver, K.M., Florez-Paz, D.M., Badea, T.C., Mentis, G.Z., Menon, V., Nooij, J.C.: Molecular correlates of muscle spindle and golgi tendon organ afferents. Nature communications 12(1), 1451 (2021).

## References

[50] Kullback, S., Leibler, R.A.: On information and sufficiency. The annals of mathematical statistics 22(1), 79–86 (1951).

[51] Pearson, K.: On the laws of inheritance in man. Biometrika 3(2-3), 131–190 (1904).

[52] Fisher, R.A.: A mathematical examination of the methods of determining the accuracy of observation by the mean error, and by the mean square error. Monthly Notices of the Royal Astronomical Society 80(8), 758–770 (1920).

[53] Brown, R.G.: Statistical forecasting for inventory control. (No Title) (1959).

[54] Vaswani, A., Shazeer, N., Parmar, N., Uszkoreit, J., Jones, L., Gomez, A.N., Kaiser, L., Polosukhin, I.: Attention is all you need. Advances in neural information processing systems 30 (2017).

[55] Pearson, K.: Vii. mathematical contributions to the theory of evolution.—iii.regression, heredity, and panmixia. Philosophical Transactions of the Royal Society of London. Series A, containing papers of a mathematical or physical character (187), 253–318 (1896).

[56] Vinyals, O., Blundell, C., Lillicrap, T., Wierstra, D., et al.: Matching networks for one shot learning. Advances in neural information processing systems 29 (2016).

[57] Haslbeck, J.M., Waldorp, L.J.: How well do network models predict observations? on the importance of predictability in network models. Behavior research methods 50(2), 853–861 (2018)

