## Supplementary Material for "DCAF: Unifying Data Validity, Evaluation Validity and Predictive Modeling for Single-cell Perturbation Simulation"

1 BGI Research, Beijing 102601, China.
2 School of Artificial Intelligence, University of Chinese Academy of Sciences, Beijing 101408, China.
3 International Centre for Bamboo and Rattan, Beijing, China.
4 BGI Research, Wuhan 102601, China.

† These authors contributed equally as the first authors.

#### Supplementary Figures

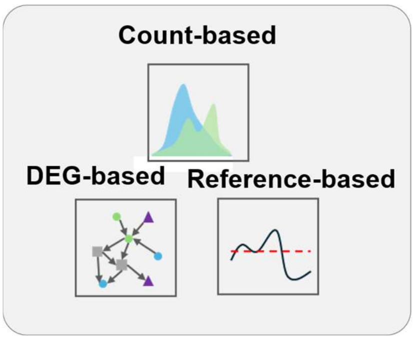


Supplementary Fig. 1: **Current criteria to assess accuracy of perturbation prediction models.**

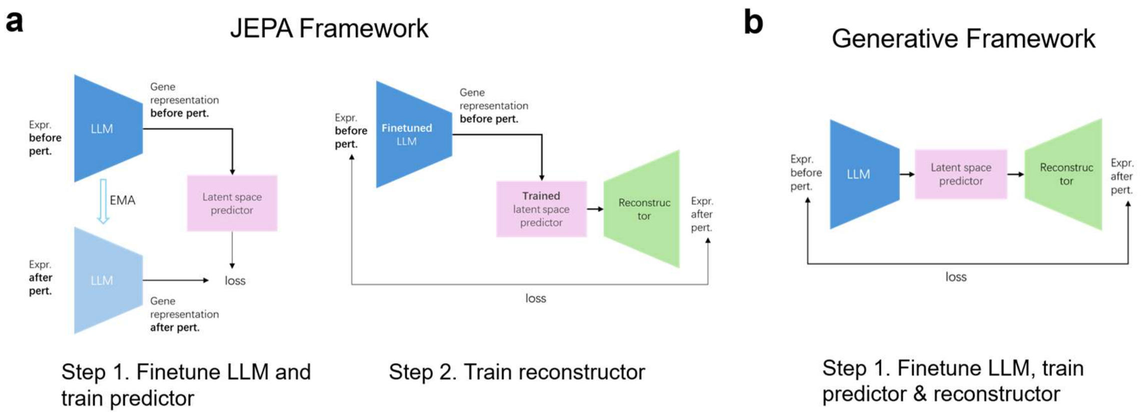


Supplementary Fig. 2: **Comparison between JEPA and custom generative framework, in terms of both general model structures and corresponding training steps.** Presented respectively in (a) and (b) are JEPA and generative frameworks.

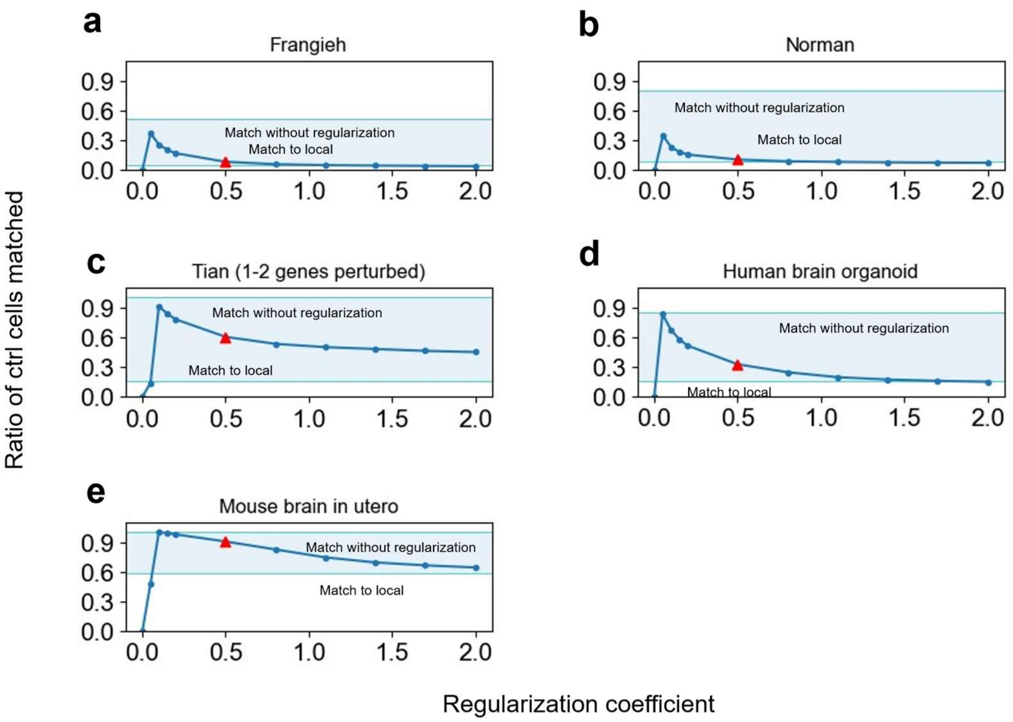


Supplementary Fig. 3: **Ratio of matched control cells caused by matching without regularization, local matching, and matching by DCAF employing varying regularization factors.** Shown in subplot (a) to subplot (e) are respectively results using dataset of Frangieh, Norman, Tian, Human brain organoid and Mouse brain *in utero*. On each subplot, upper arrow points to result by matching without regularization and lower arrow points to result by locally restricted matching. Dark blue dots denote ratio caused by changing regularization factor in DCAF matching, and the trend is shown by dark blue lines. Red triangle highlights ratio of matched control cells caused by the chosen regularization parameter.

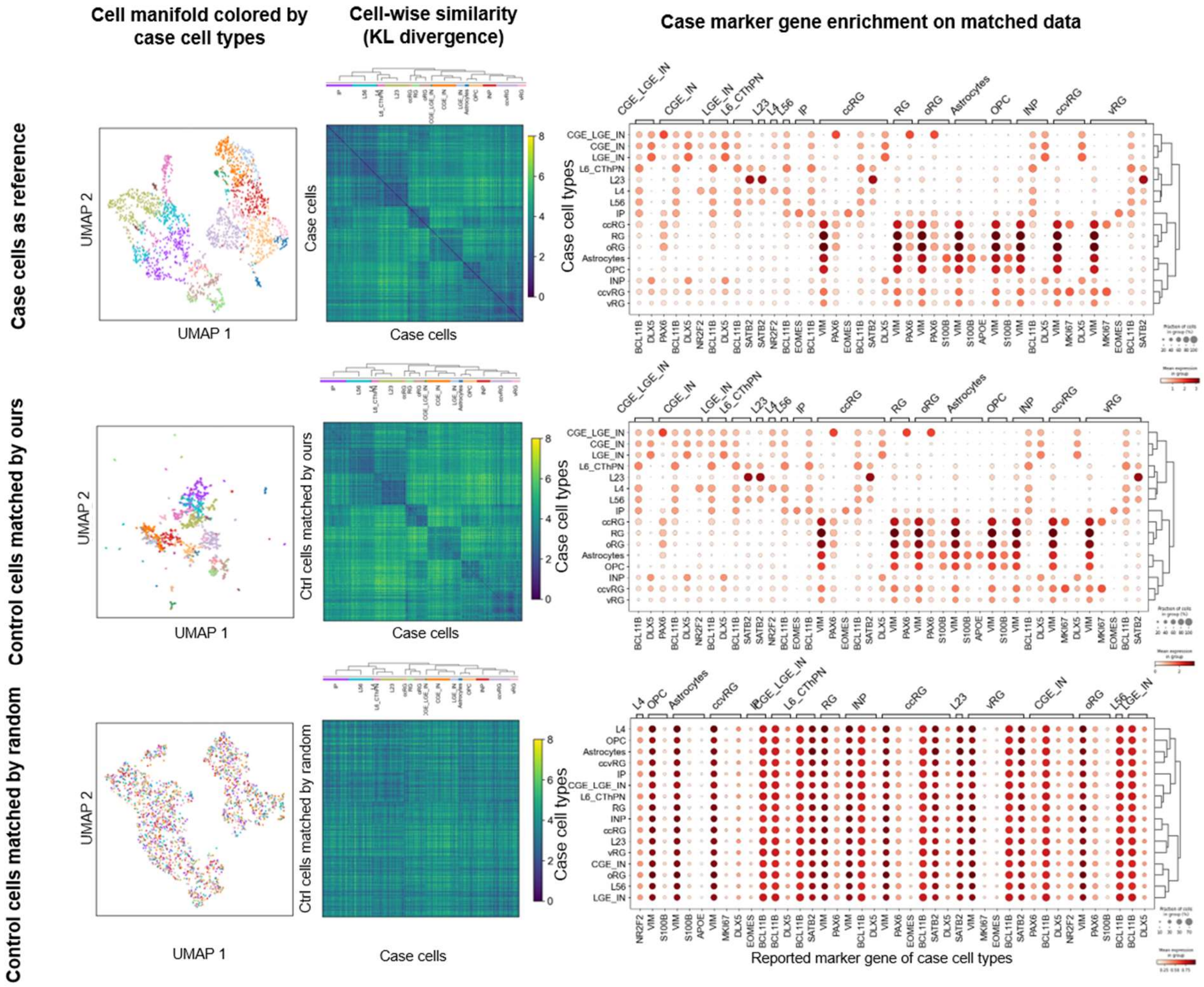


Supplementary Fig. 4: **Comparison of intrinsic cell identity similarity and marker gene information on Human brain organoid recovered by different ways.** Present in the 1st column are UMAP colored by matched case cell type, in the 2nd column are KL divergence heatmaps, and in the 3rd column are marker enrichment plots. From top to bottom presents recovered information from different sources, ranging from case cells themselves, control cells matched by DCAF, to randomly matched control cells.

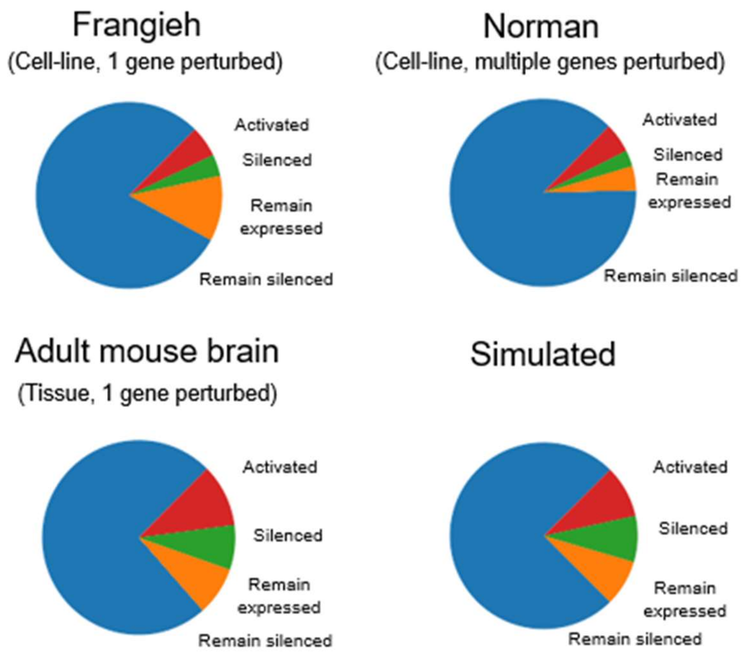


Supplementary Fig. 5: **Gene expression changes composition from cell line, multi cell type datasets, and simulated dataset.**

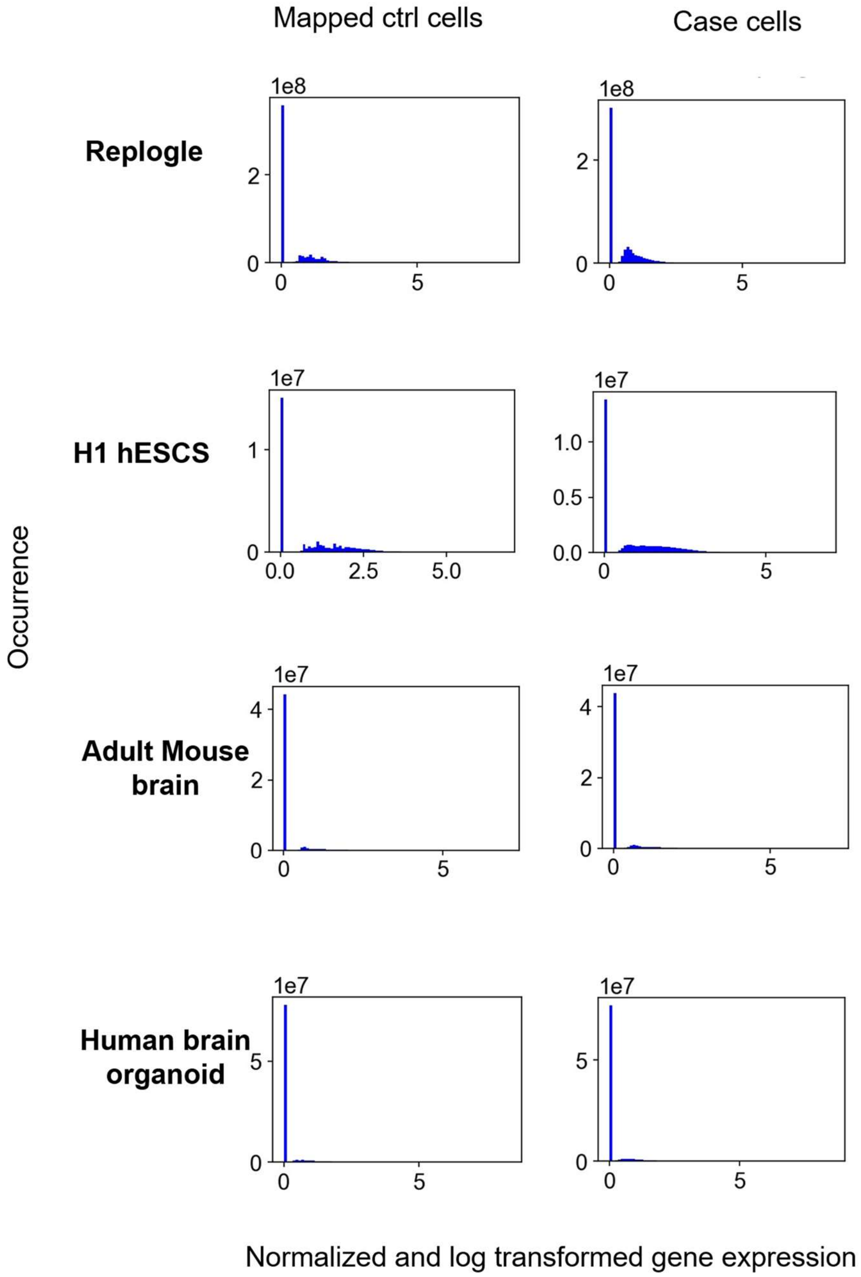


Supplementary Fig. 6: **Distribution of normalized and log transformed gene expression of control cells matched by DCAF, and case cells.** Present from the 1st to the 4th row are datasets of Replogle, H1 hESCS, adult Mouse brain and Human brain organoid. Present on left are expression distributions of mapped control cells and in right are distributions of case cells.

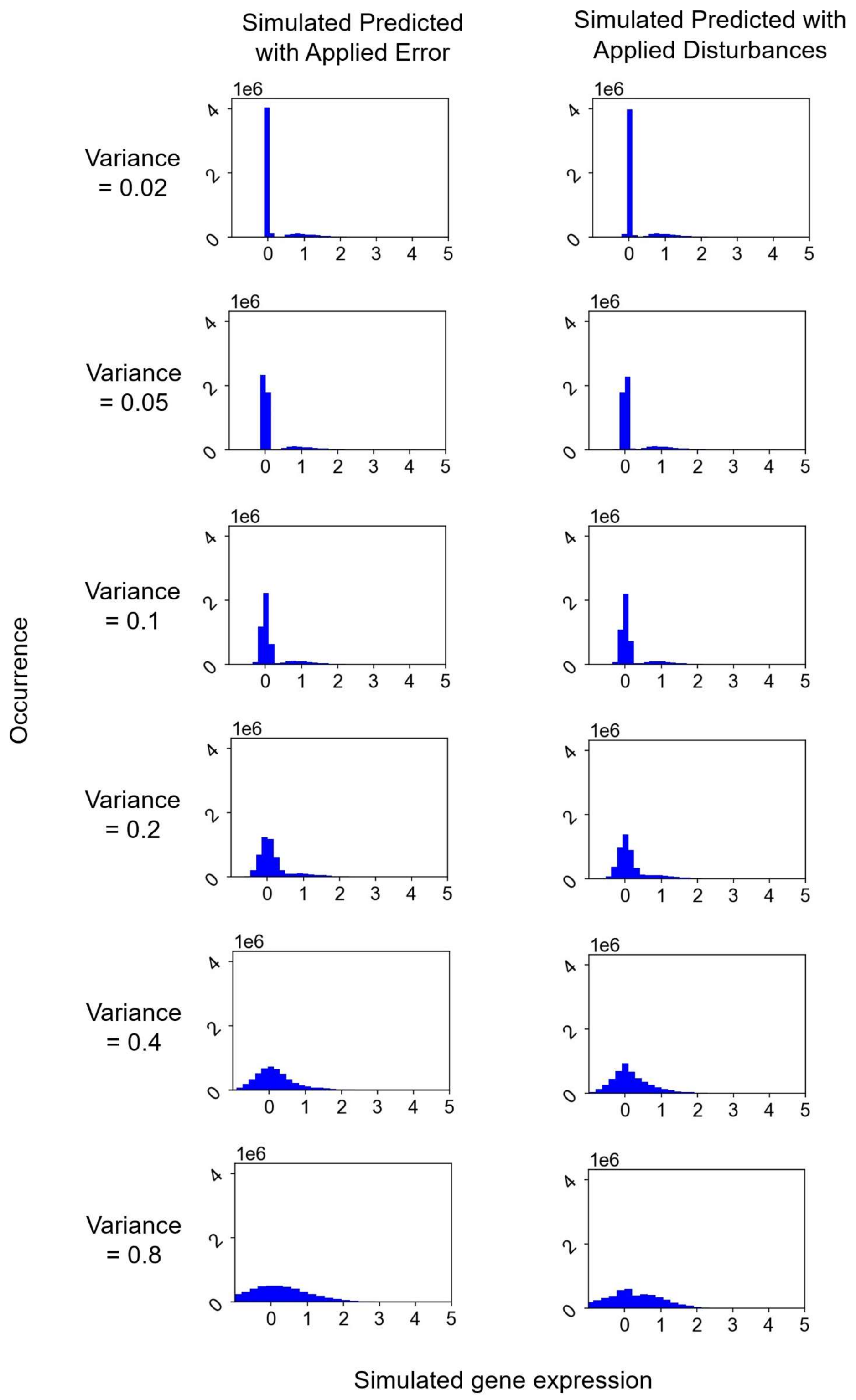


Supplementary Fig. 7: **Simulated prediction datasets with known levels of error and disturbance.** Present on left are distributions of simulated predicted expression with applied error, and in right are distributions of simulated predicted expression with applied disturbances. From top to bottom are results by applied variance of 0.02, 0.05, 0.1, 0.2, 0.4 and 0.8.

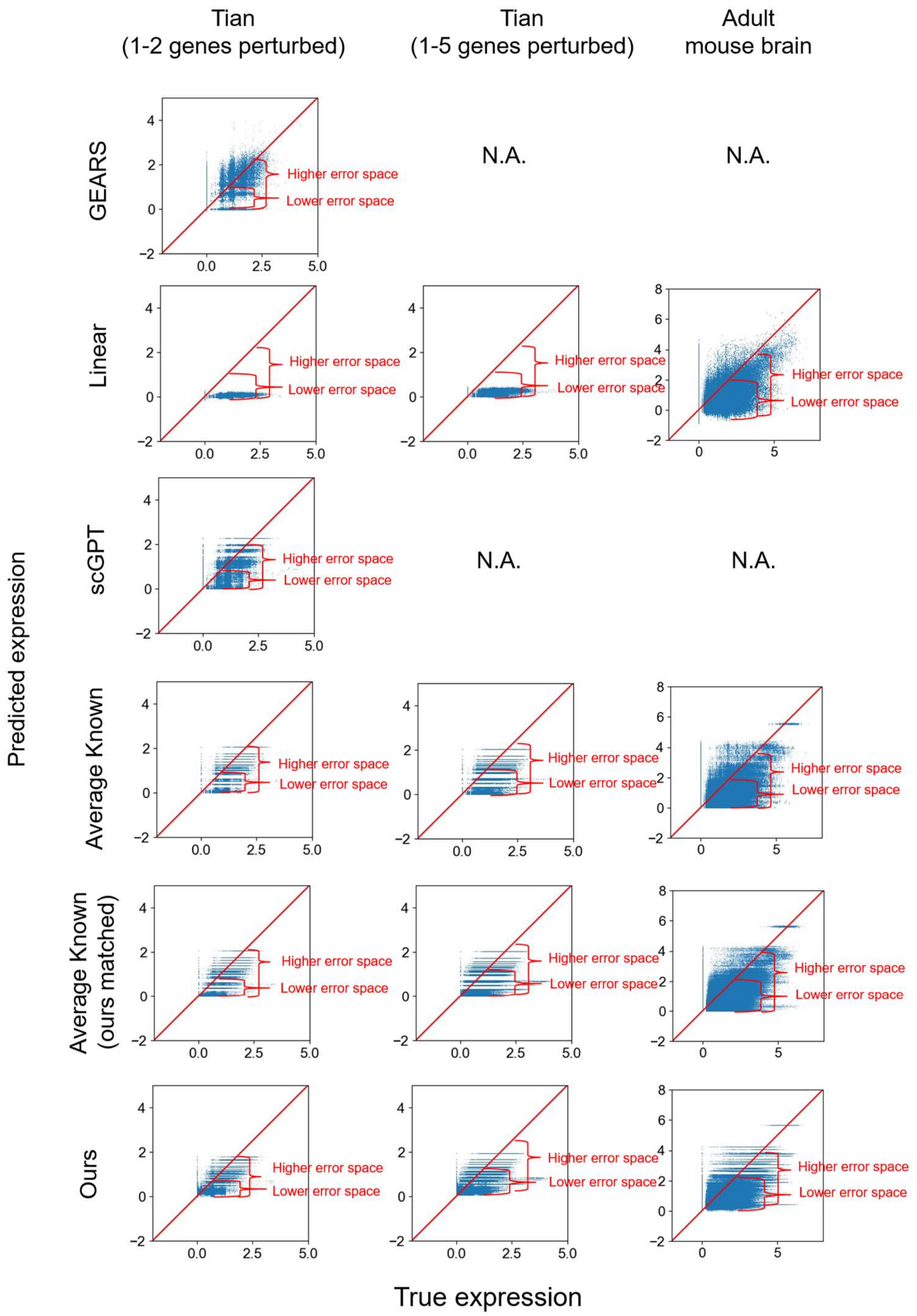


Supplementary Fig. 8: **Comparison between predicted, as well as true normalized then log transformed gene expression across different methods on different datasets.** Present to top to bottom are results by GEARS, Linear, scGPT, Average Known and ours. Present from left to right are results of datasets of Tian (1-2 genes perturbed), Tian (1-5 genes perturbed) and adult Mouse brain. Strict alignment is marked by red solid lines and comparison of error space between different true expression levels are marked by red parenthetical annotations.

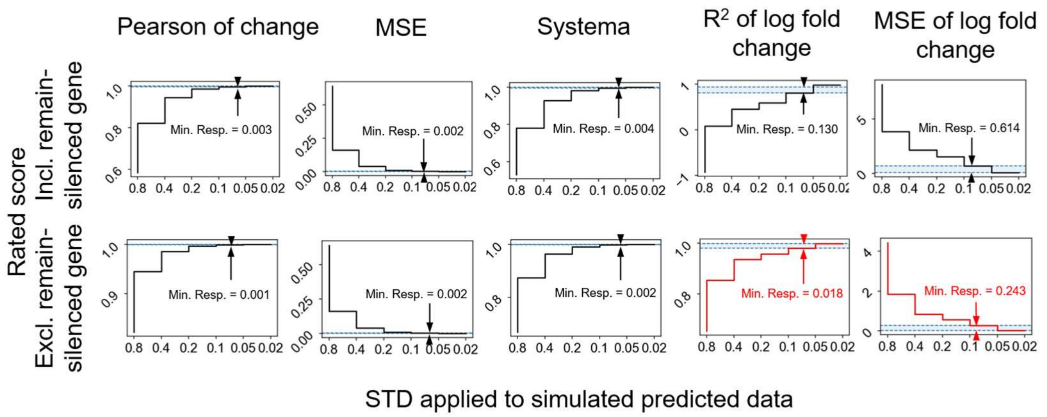


Supplementary Fig. 9: **Comparison of different criteria's sensitivity between cases when remain-silenced genes are included and excluded.** Present on the 1st and 2nd rows are rated scores of criteria in response to different STD marked on x axes, applied to simulated predicted data, respectively when remain-silenced genes are included and excluded for each cell. Shown from left to right are respectively criteria including Pearson of change, MSE, Systema, $R^2$ of log fold change and MSE of log fold change. Marked by arrow annotations are minimum change of rated scores in response to exponentially decreasing STD. Visualized by red are performance of DCAF criteria.

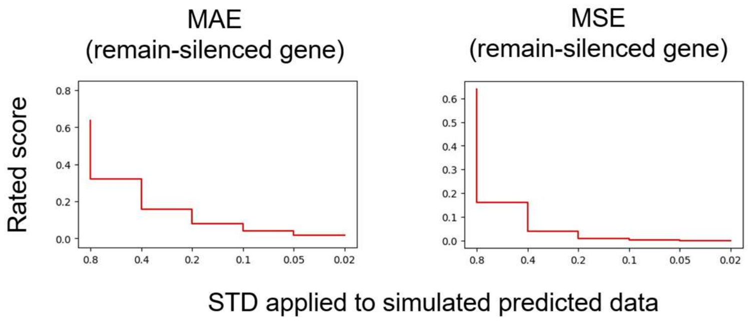


Supplementary Fig. 10: **Response of DCAF's criteria on remain-silenced gene to varying STD applied to simulated predicted data.** Present in the 1st column is MAE of expression while present in the 2nd column is MSE of expression.

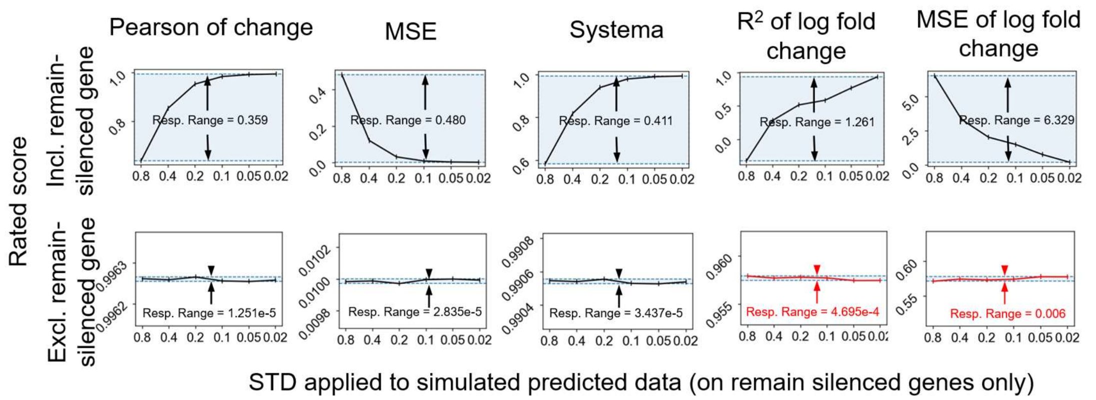


Supplementary Fig. 11: **Comparison of different criteria's robustness between cases when remain-silenced genes are included and excluded.** Present on the 1st and 2nd rows are rated scores of criteria in response to different STD marked on x axes applied to remain silenced genes of simulated predicted data, respectively when remain-silenced genes are included and excluded for each cell. Shown from left to right are respectively criteria including Pearson of change, MSE, Systema, $R^2$ of log fold change and MSE of log fold change. Marked by arrow annotations are response range of rated scores in response to exponentially decreasing STD. Visualized by red are performance of DCAF criteria.

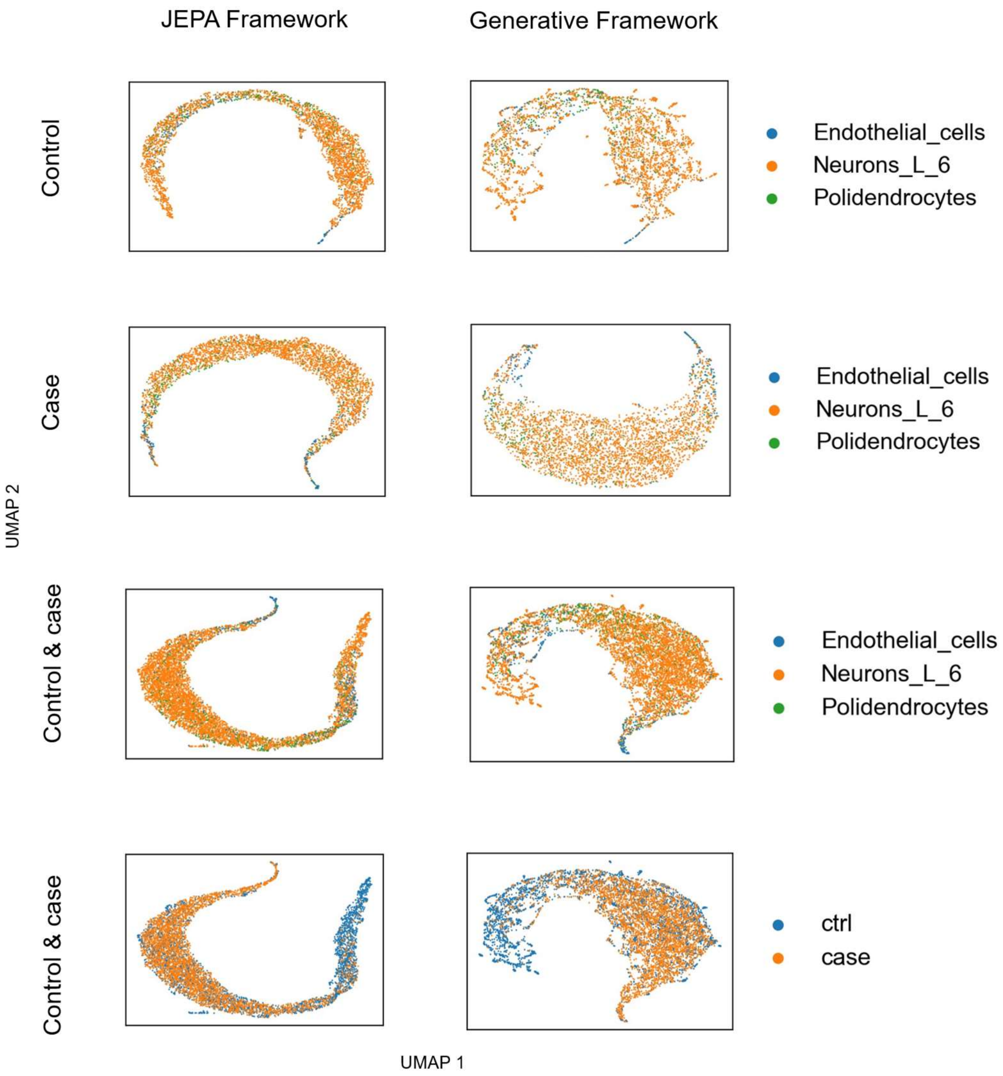


Supplementary Fig. 12: **Comparison of cell embeddings generated by JEPA versus generative framework using adult Mouse brain dataset.** Present from left to right are results of JEPA and generative frameworks. Present from top to bottom are respectively UMAP of control cell embeddings colored by cell types, case cell embeddings colored by cell types, control and case cells together colored by cell types, and control and case cells together colored by perturbation status.

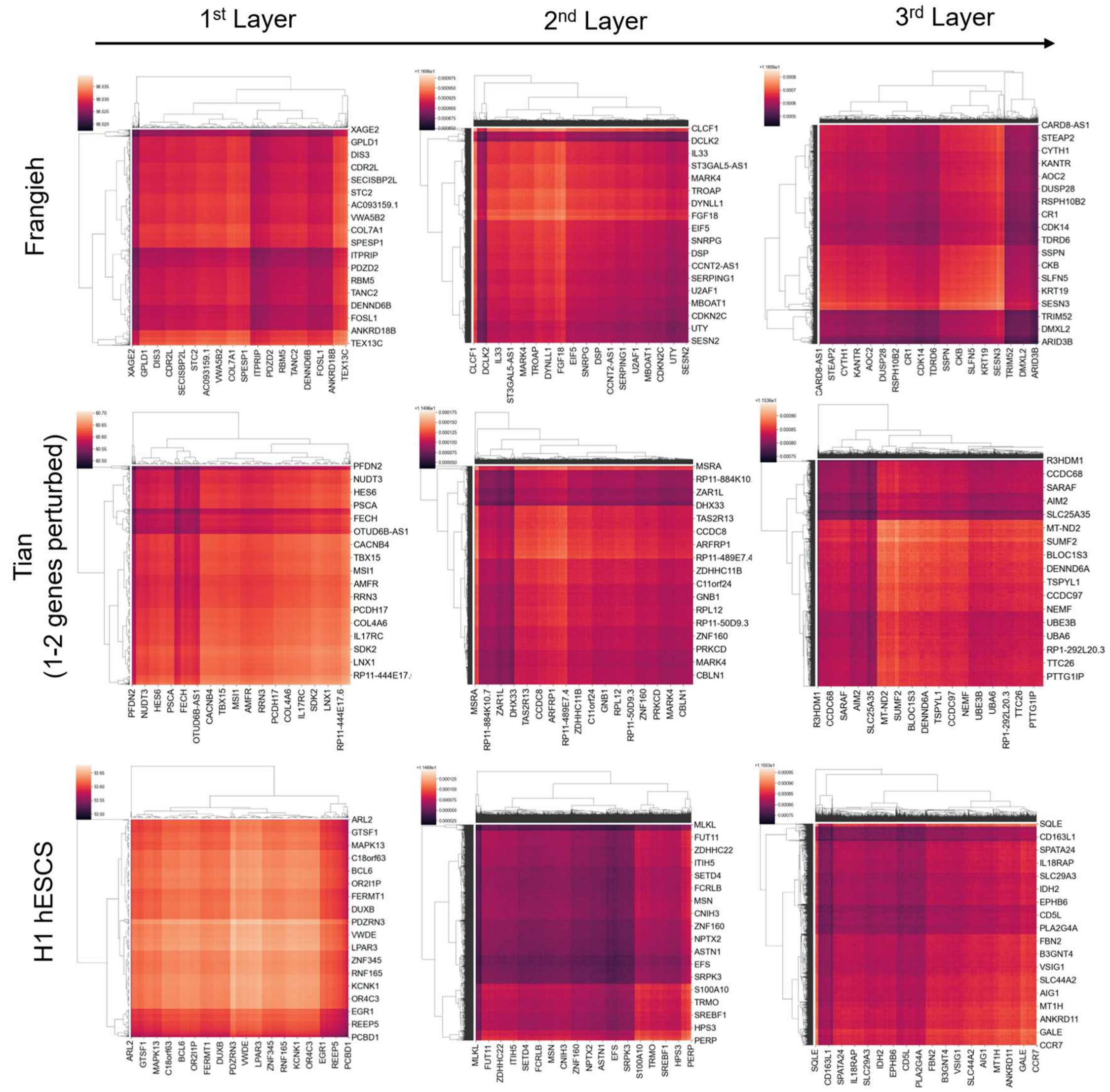


Supplementary Fig. 13: **Clustered attention maps generated from the 1st to 3rd layer of our predictor, using datasets of Frangieh, Tian (1-2 genes perturbed) and H1 hESCS.** Presented from left to right are maps generated by 1st, 2nd and 3rd layers. Shown from top to bottom are respectively results of Frangieh, Tian (1-2 genes perturbed) and H1 hESCS datasets. A testing cell not seen by model is used for visualization of each dataset.

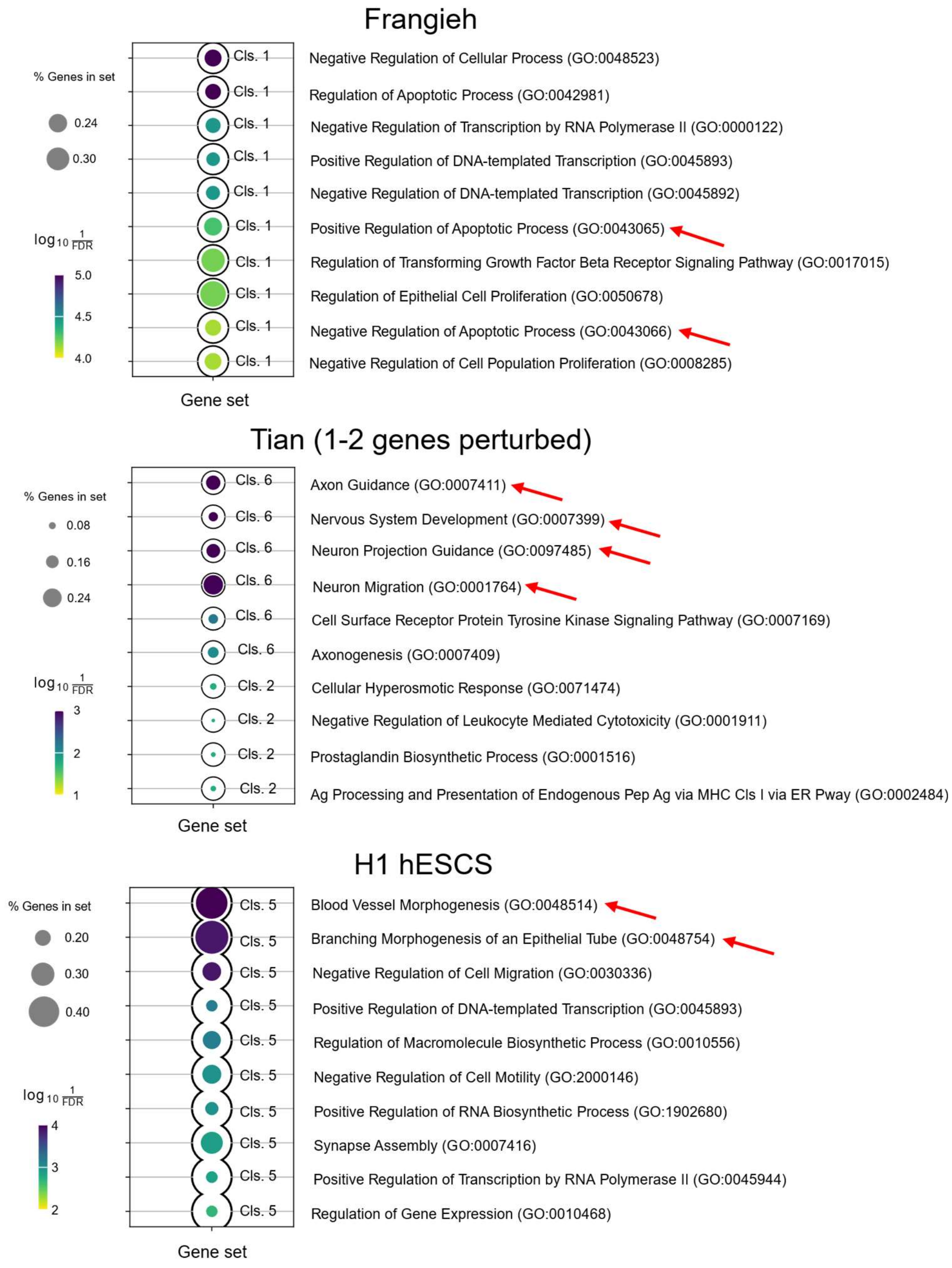


Supplementary Fig. 14: **Gene enrichment analysis for gene clusters generated by attention maps by the 3$^{rd}$ layer of our predictor, using datasets of Frangieh, Tian (1-2 genes perturbed) and H1 hESCS.** Shown from top to bottom are respectively results of Frangieh, Tian (1-2 genes perturbed) and H1 hESCS datasets. Apoptotic functions enriched for Frangieh dataset measuring Human lung cancer, neuronal functions enriched for Tian dataset measuring developing neurons, and morphogenesis functions enriched for H1 hESCS measuring brain organoid are pointed by red arrows.

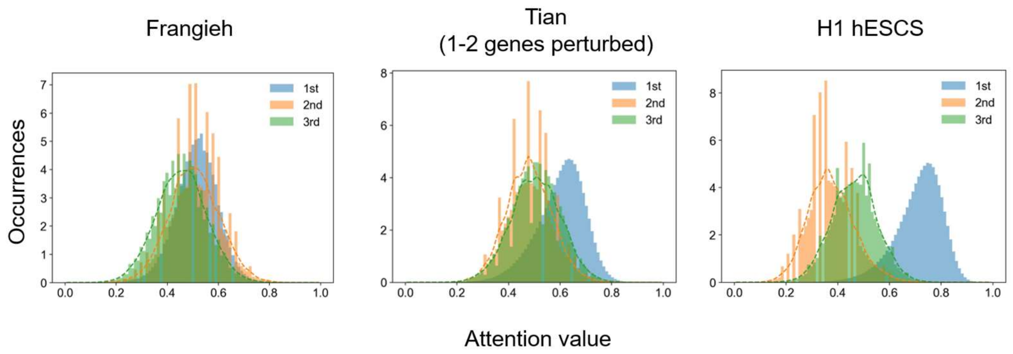


Supplementary Fig. 15: **Distribution of min-max scaled attention values generated from the 1st, 2nd and 3rd layer of our predictor, using datasets of Frangieh, Tian (1-2 genes perturbed) and H1 hESCS.** Shown from left to right are respectively results of Frangieh, Tian (1-2 genes perturbed) and H1 hESCS datasets. Gaussian denoising results of the 1st and 2nd layers are respectively highlighted using orange and green dashed lines.

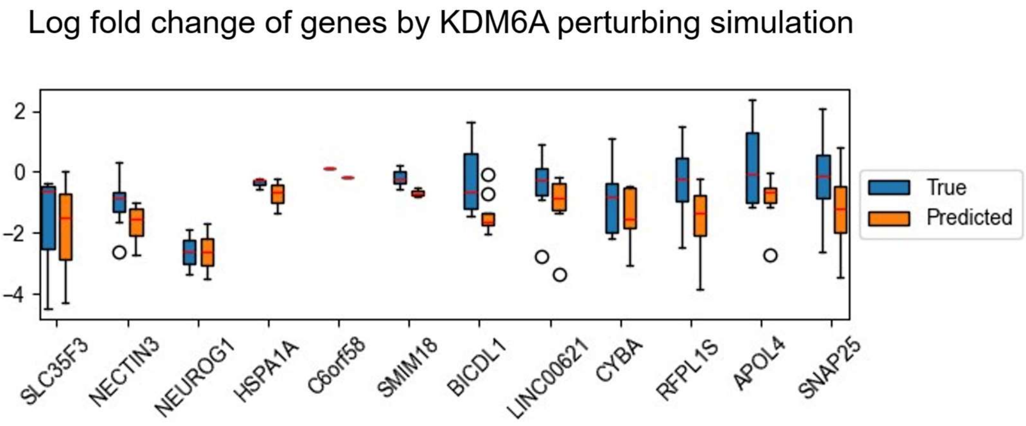


Supplementary Fig. 16: **Comparison between predicted and true log fold change across highly variable genes by KDM6A perturbation using Human brain organoid dataset.** The whiskers extend from the box to the farthest data point lying within 1.5 times the inter-quartile range (IQR) from the box. Median values of each data are highlighted by red solid lines.

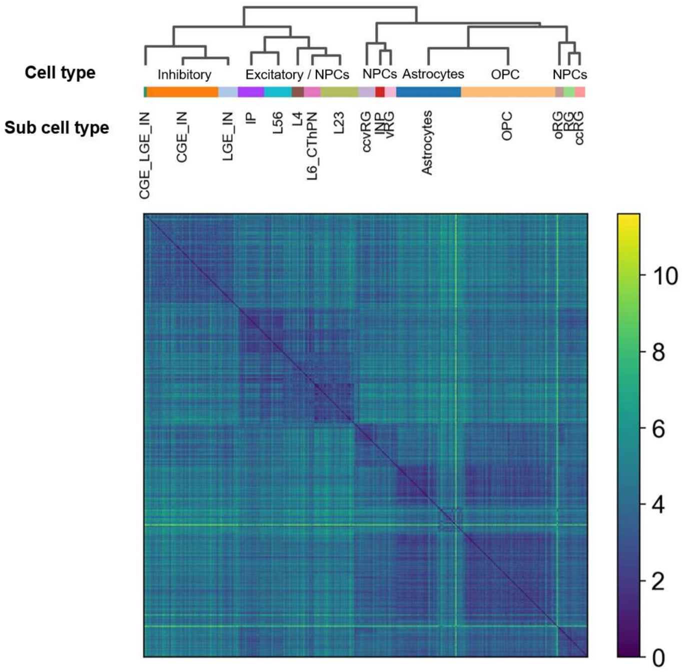


Supplementary Fig. 17: **Single-cell level KL divergence between clustered then annotated cells using Human brain organoid dataset.**

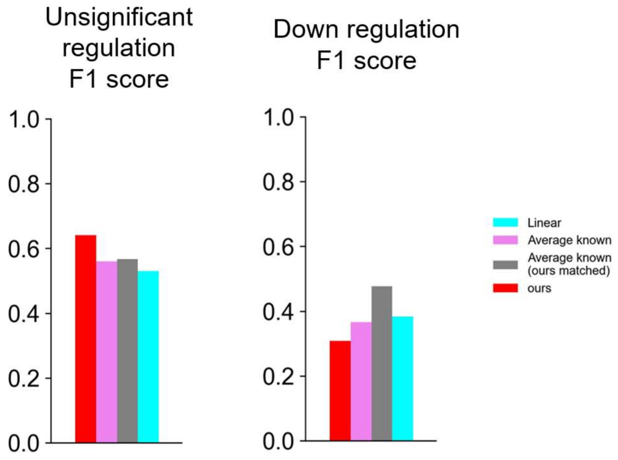


Supplementary Fig. 18: **Unsignificant regulation and down regulation F1 scores of different methods including Linear, Average known and ours, using adult Mouse brain dataset.** Present on the left are unsignificant regulation F1 scores and present on the right are down regulation F1 scores.

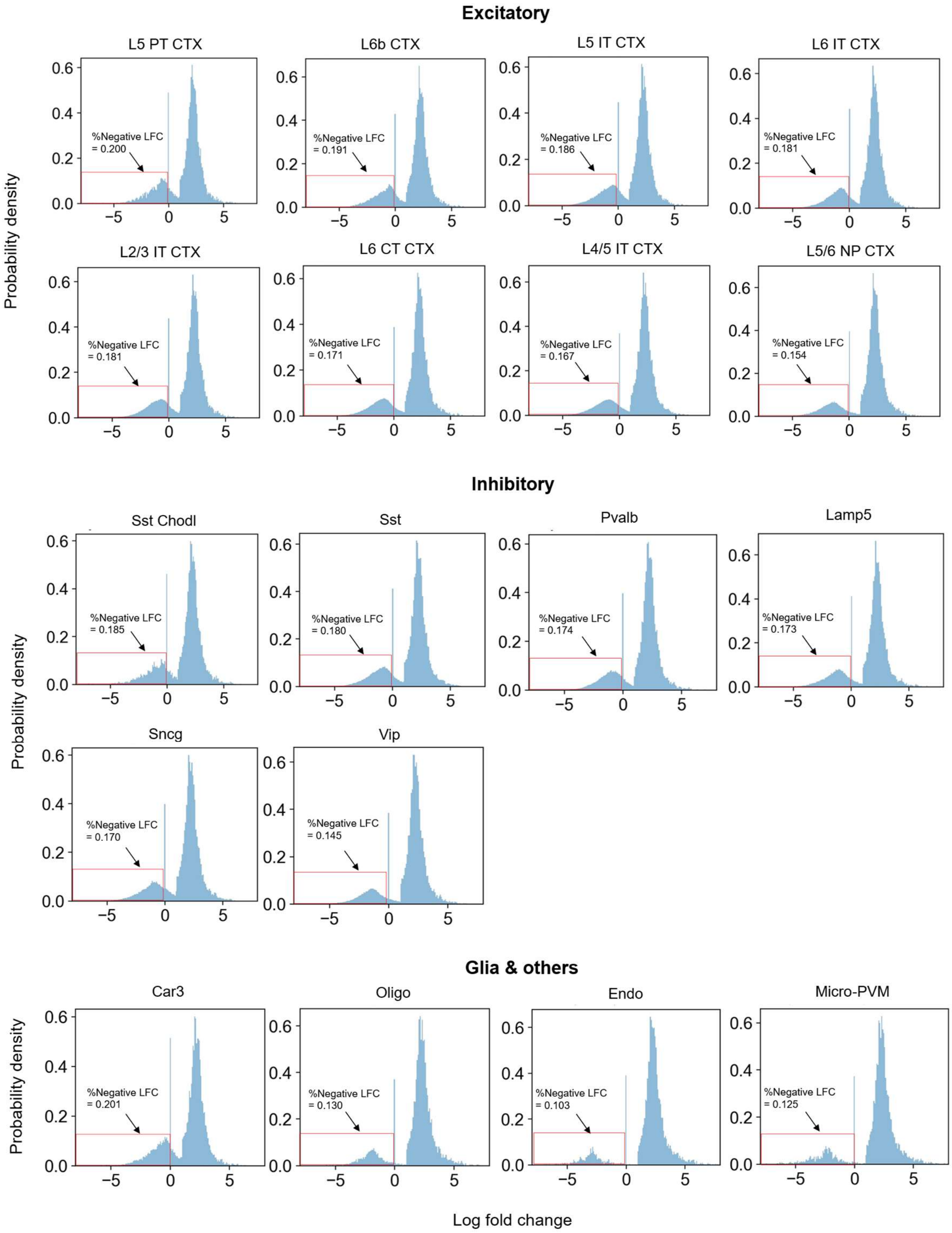


Supplementary Fig. 19: **Distribution of log fold change across different genes of different cells for excitatory, inhibitory and glial/other neurons.** Negative log fold change part is squared by red solid lines, with their ratios highlighted by arrow annotations.

### Supplementary Tables

Supplementary Table 1: **Summary of no. of cells that are perturbed and not perturbed for adopted datasets.**

| Dataset | No. of control cells | No. of case cells | Ratio of control over case cell no. | (Match to local) ratio of matched control cells |
|---|---|---|---|---|
| Adult mouse brain | 2520 | 12958 | 0.194 | 0.398 |
| Tian (1-5 genes perturbed) | 2477 | 24875 | 0.100 | 0.294 |
| Frangieh | 18334 | 11142 | 1.645 | 0.051 |
| Norman | 11855 | 11300 | 1.049 | 0.081 |
| Tian (1-2 genes perturbed) | 2477 | 6333 | 0.391 | 0.157 |
| Human brain organoid | 14703 | 222863 | 0.643 | 0.160 |
| Mouse brain *in utero* | 374 | 11212 | 0.033 | 0.591 |

Supplementary Table 2: **Summary of low threshold and DCAF accuracy for remaining silenced genes of adopted datasets.**

| Dataset | Low threshold | MAE (of genes that remain silence) | MSE (of genes that remain silence) |
|---|---|---|---|
| H1 hESCS | 0.1 | 0.267 | 0.123 |
| Norman | 0.1 | 0.2 | 0.068 |
| Frangieh | 0.1 | 0.283 | 0.110 |
| Tian (1-2 genes perturbed) | 0.05 | 0.216 | 0.069 |
| Replogle | 0.1 | 0.372 | 0.158 |
| Tian (1-5 genes perturbed) | 0.05 | 0.256 | 0.075 |
| Human brain organoid | 0.1 | 0.254 | 0.094 |
| Adult mouse brain | 0.1 | 0.356 | 0.171 |
| Mouse brain *in utero* | 0.1 | 0.569 | 0.718 |
